# *FBXW11* Activity Regulates Radial Glial Expansion in Human Cerebral Organoids

**DOI:** 10.64898/2026.08.20.746070

**Authors:** Cesar L Moreno, Helen King, Sophia Trabish, Marloes Thijs, Dominik Beck, Maria Bergamasco, Tian Y. D’Araujo, Timothy J. Mosca, Rob Weatheritt, G. Gregory Neely

## Abstract

Human brain development depends on tightly coordinated gene-regulatory programs and the emergence of complex tissue architecture, making large scale functional interrogation difficult using conventional screen models. To overcome this challenge, we used a pooled CRISPR screening approach. Guided by neuro-specific whole-genome screens in *Drosophila*, we tested 129 poorly characterised human orthologs and found 8 that modify cerebral organoid development. Candidates were validated using individual CRISPR knockouts and mosaic competition assays. Among these candidates we describe *FBXW11*, a substrate-recognition component of the SCF E3 ubiquitin ligase complex, as a potent negative regulator of cerebral organoid expansion. *FBXW11* loss increases radial glial abundance, expands ventricular-like domains, and impairs neuronal maturation. Mechanistically, FBXW11 associates with β-catenin and alters WNT signalling. *FBXW11* mutations cause the autosomal-dominant Mendelian syndrome *Neurodevelopmental, Jaw, Eye and Digital syndrome* (NEDJED), and we found that disease-associated variants mapped preferentially to WD40 substrate-binding repeats and β-catenin contact regions, linking impaired substrate recognition to neurodevelopmental disease. Together, these findings identify *FBXW11* as a conserved negative regulator of β-catenin-dependent radial glial expansion and neuronal maturation during human cerebral brain development.

## Main

The development of the human brain is driven by highly orchestrated gene regulatory programs that coordinate progenitor expansion, neuronal differentiation, migration, and tissue architecture^1^. Identifying new regulators of these processes remains an active area of developmental research and is essential for understanding both normal brain function and disease. Human genetic studies have revealed many disease-associated loci, but genes without established links to Mendelian disease, GWAS signals, or recurrent *de novo* variation are less frequently prioritized for functional studies, even though they may encode conserved regulators of brain development.

Comparative genetics provides an alternate route to uncover such biology. Model organisms such as *Drosophila* have enabled systematic gene discovery at genome scale, broadly cataloguing genes required for nervous system development^2-5^. Importantly, many of the pathways identified in these invertebrate models show strong conservation of function across phyla, highlighting how comparative genomics can provide insight into human biology^6^. More recently, stem cell-derived cerebral organoids have emerged as powerful models of human neurodevelopment, enabling investigation of developmental programs that cannot be readily studied in animal models while providing mechanistic insight into neurological disease^7^.

To identify previously underexplored regulators of human brain development we applied a pooled CRISPR screening approach^8^. We curated genome-wide nervous system lethality datasets generated in *Drosophila* using the pan-neuronal driver *elav-Gal4* and the neural stem cell driver *inscuteable-Gal4*^4,5^. We then prioritised evolutionarily conserved genes with limited functional annotation, reasoning that this strategy could reveal conserved developmental regulators not yet interrogated in human cerebral organoids. We evaluated these candidates using a barcode-enabled pooled CRISPR screening platform for human cerebral organoids. From this, we identified 8 new regulators of cerebral organoid development, including *FBXW11* (F-Box and WD Repeat Domain Containing 11) which we show is a regulator of organoid growth, cellular composition, cell-cycle state, and neuronal maturation Mechanistically, FBXW11 interacts with β-catenin to restrain WNT signalling, supporting a model where FBXW11 limits proliferating radial glial cells during human cerebral organoid development.

## Results

### A barcoded pooled CRISPR screen in cerebral organoids identifies new regulators of neurodevelopment

To identify regulators of human cortical development we designed a CRISPR knockout (KO) library from genome-wide *Drosophila* nervous-system lethality datasets. Specifically, we began with a set of fly genes whose knockdown by the pan-neuronal driver *elav-*Gal4 or the neural-stem-cell driver *inscuteable-*Gal4 produced lethality^4,5^, returning 1,677 genes with a 9.4% overlap (158 genes) across drivers. To enrich for conserved neurodevelopmental genes with limited prior annotation in human cortical development, we applied two complementary filtering strategies (Fig.1a). In the first, we filtered fly genes by Flybase Gene Ontology assignments of unknown / unannotated function^9^, then retained those with a high-confidence human ortholog (>10 DIOPT score^10^), yielding 99 genes. In the second, we first identified high-confidence human orthologues, then enriched for human GO-unannotated terms^11^, yielding 68 genes. Combining these pipelines produced a final 129-gene library, together with positive and negative controls (Extended Data Table 1). We hypothesized that this approach would maximize discovery of gene regulators that combined (1) likely functionality during the nervous system development (based on their reported invertebrate lethality), and (2) unknown or modest information on their role in brain development.

Using these candidates, we constructed a CRISPR-KO library using 4 guides/gene (Fig. 1b), together with positive and intergenic negative control guides. Recent studies show that pooled CRISPR-screening when performed in heterogenous tissues, such as cerebral organoids, can be confounded by stochastic clonal expansion leading to false target identification^8^. We therefore incorporated an additional nucleotide barcode within the guide vector to support barcode-level interpretation of enrichment and depletion (Extended Data Fig.1a). We termed this design ORIGAMI, for Organoid Resolved Interrogation of Genetic Activity using Molecular Identifiers (see methods). Sequencing results of the plasmid library confirmed high representation and diversity (Gini index = 0.08), supporting that we could proceed to screen with comprehensive coverage. To evaluate whether this barcode layer could reduce founder-driven variability, we analysed intergenic negative-control guides, which should remain neutral during differentiation. Barcode recovery was high across all replicates and time points, enabling barcode-level quality control throughout the screen (Extended Data Fig. 1b). Raw barcode counts showed increasing mean–variance dispersion as organoids differentiated, consistent with stochastic expansion of individual barcoded clones in three-dimensional culture (Extended Data Fig. 1c). After outlier discrimination, barcode count distributions showed reduced inequality (Gini index) and improved Simpson diversity, with the largest effects observed after transition into organoid culture rather than at DIV0 (Extended Data Fig. 1d,e). These data agree with previously identified artifacts in 3D organoid screens^8^.

**Fig. 1.**
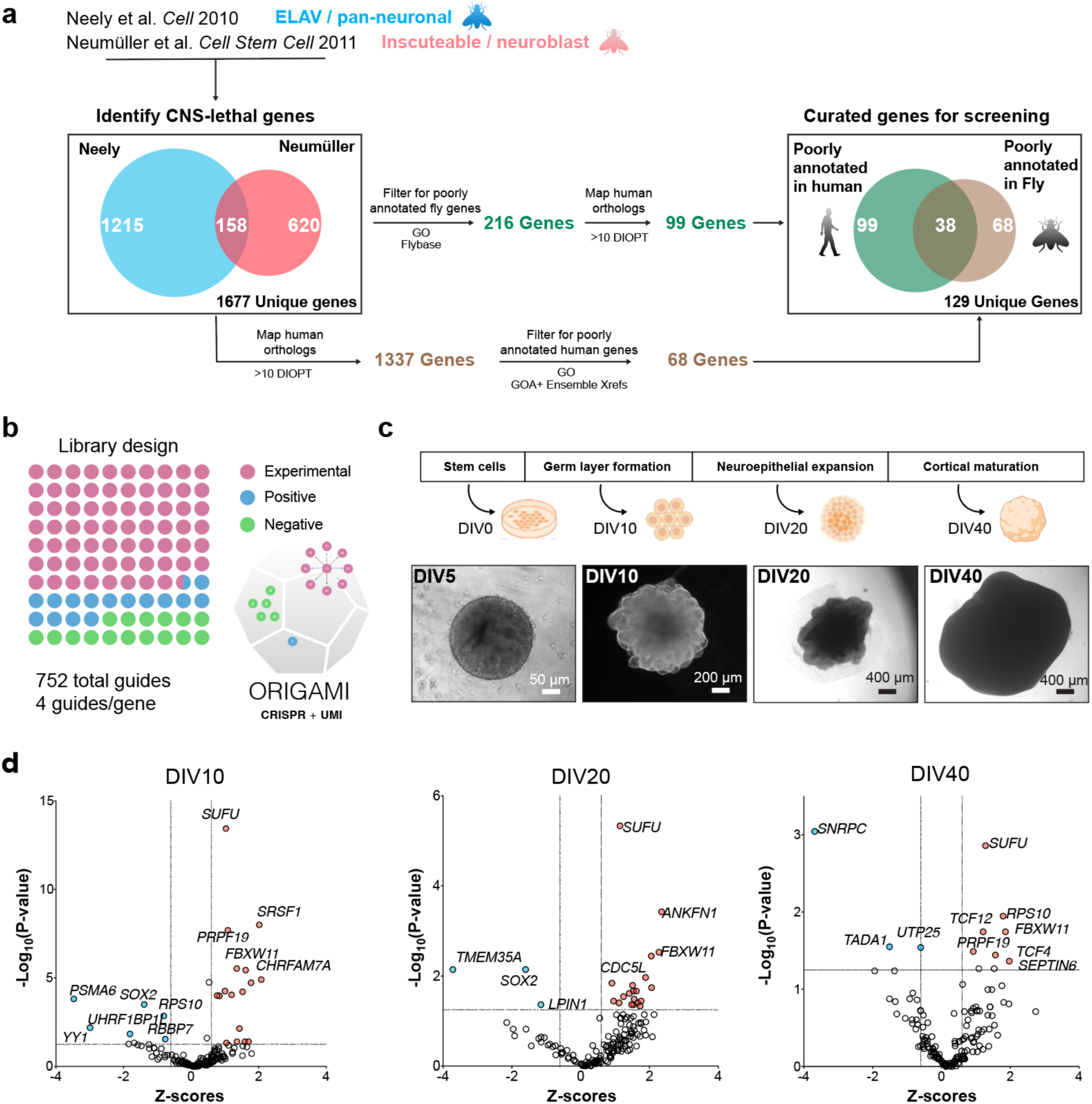
ORIGAMI enables barcoded pooled CRISPR screening in human cerebral organoids. **a,** Schematic of the comparative genomics pipeline used to identify understudied genes with conserved nervous-system relevance. **b,** Schematic of the library design. **c,** Experimental workflow and sampling timepoints for pooled CRISPR screening during cerebral organoid differentiation. **d,** Volcano plots showing enriched and depleted gene targets at DIV10, DIV20, and DIV40 of cerebral organoid differentiation. N = 3 experimental replicates.

Since we first identified our genes of interest from Fly screens restricted to the nervous system, we used a cerebral brain organoid protocol, over more specific directed protocols, to capture broader neurodevelopmental effects^12^. We focused on early timepoints of cerebral organoid brain differentiation (Fig. 1c), and detected enrichment and drop out of targets across day 10, 20, and 40 of differentiation (Fig.1d). Across all independent experimental replicates and timepoints, sgRNA with barcodes were uniformly detected across all groups (Extended Data Fig. 1b). As expected, *SOX2*, a key regulator of pluripotency and neural progenitor identity, was depleted at early stages, whereas KO of the brain tumour suppressor *SUFU* was enriched across differentiation. Additional positive-control and cell-fitness genes including *TOP1* (encoding for DNA topoisomerase I), *FGFR1* (encoding for a fibroblast growth receptor), *RPL8* (Ribosomal protein), *NUMB* (a gene involved in asymmetrical cell division), and *VENTX* (a homeobox gene), were also under-represented at baseline or early time points (Extended Data Fig. 1d). Together, these results show that ORIGAMI can detect gene-dependent changes in cellular fitness and developmental growth across early cerebral organoids.

### Orthogonal validation identifies candidate regulators of cerebral organoid size

To validate screen hits, we generated pooled iPSC lines individually targeted with single sgRNAs against selected candidate genes. We hypothesised that KOs identified as either enriched or depleted would produce larger or smaller cerebral organoids, respectively (Fig. 2a). Targeted genomic regions were PCR amplified from each cell edited pool, and KO efficiency was estimated by Sanger trace analysis using Inference of CRISPR Edits (ICE)^13^ (Fig. 2b). We found that most guides resulted in at least a ∼50% gene disruption, except for *CDC5L*. We then differentiated each edited pool into cerebral organoids and quantified organoid area at DIV40. Individual targeting reproduced size phenotypes for substantial fraction of candidates, with significant differences observed in 60% of genes tested (Fig. 2c, d). Among these, disruptions of Transcriptional Adaptor 1 (*TADA1)* produced the smallest organoids, whereas loss of F-box/WD repeat-containing protein 11 (*FBXW11)* produced the largest.

**Fig. 2.**
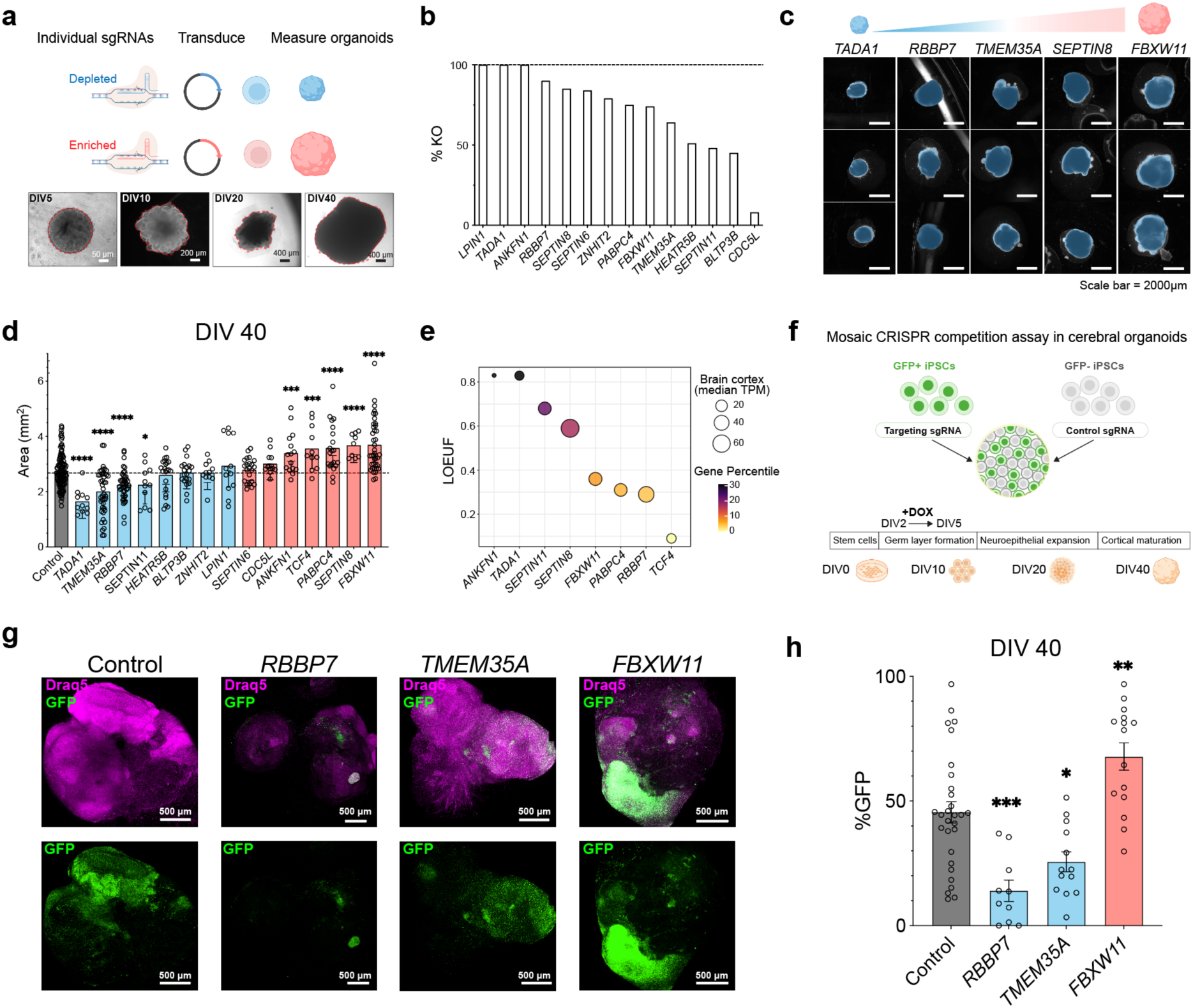
Validation of candidate regulators identifies genes that increase or decrease cerebral organoid growth. **a,** Workflow for secondary validation of candidate genes identified from the ORIGAMI screen. Individual sgRNAs targeting candidate genes predicted to decrease or increase organoid growth were used to generate edited iPSC pools, which were differentiated into cerebral organoids and quantified at DIV40. **b,** Editing efficiencies determined by targeted amplicon sequencing and ICE analysis ^13^. **c,** Representative brightfield images of DIV40 cerebral organoids following candidate gene targeting. **d,** Quantification of organoid area at DIV40 following CRISPR-mediated knockout. Each dot represents an individual organoid across independent differentiations. Bars indicate mean ± SEM, n > 3 independent differentiations. Statistical significance was determined relative to control using one-way ANOVA; \**P* < 0.05, \*\*\**P* < 0.001, \*\*\*\**P* < 0.0001. **e,** Human cortical expression and genetic constraint of validated targets. Dot size and colour indicate median cortical expression (TPM) and gene ranking percentile, respectively, while y-axis shows loss-of-function intolerance (LOEUF). **f,** Schematic of the mosaic CRISPR competition assay used to evaluate candidate gene effects on organoid fitness. DOX-inducible Cas9 iPSCs expressing eGFP-NLS were transduced with candidate sgRNAs, mixed with GFP-negative control cells, and differentiated into cerebral organoids. Cas9 expression was induced between DIV2 and DIV5 and GFP-positive cells were quantified at DIV40. **g,** Representative images of mosaic cerebral organoids at DIV40. GFP-positive cells carrying sgRNAs targeting *RBBP7*, *TMEM35A*, or *FBXW11* were competed against control cells within the same organoid. Draq5 labels nuclei. **h,** Quantification of the percentage of GFP-positive cells within mosaic organoids at DIV40. Data are presented as mean ± SEM, n = 3 independent differentiations. Statistical significance was determined relative to control using one-way ANOVA; \**P* < 0.05, \*\**P* < 0.01, \*\*\**P* < 0.001.

Because multiple validated candidates altered organoid size, we next integrated phenotypic results with human cortical expression and genetic constraint. Candidate genes were compared using median cortical expression^14^ and loss-of-function intolerance^15^ (Fig. 2e), reasoning that highly expressed and constrained genes would be more likely to have biologically relevant roles in human neurodevelopment. *FBXW11* and *RBBP7* combined robust cortical expression with intolerance to loss-of-function variation, whereas candidates such as *TADA1* and *ANKFN1* showed stronger constraint but lower cortical expression. Based on these analyses and the magnitude of their organoid phenotypes, we selected *RBBP7* and *FBXW11* for further validation because they represented contrasting microcephalic and macrocephalic outcomes. We also included *TMEM35A* because it showed substantial cortical expression (25.49 median TPM), a strong microcephalic phenotype (Fig. 2d), and limited available constraint information.

We tested candidate effects in a mosaic competition format, by generating a nuclear GFP-expressing iPSC line together with an isogenic GFP-negative control population. GFP-positive cells were targeted with candidate sgRNAs using a DOX-inducible CRISPR KO strategy, whereas GFP-negative cells carried control sgRNAs (Fig. 2f). Note, that we used a different background iPSC line for these studies, supporting that the phenotypes observed are general and not line dependent. These populations were mixed during cerebral organoid generation, and KO was induced by DOX administration for three days beginning at day 2. At DIV40, we quantified the proportion of GFP-positive cells within mosaic organoids. Targeting RBBP7 and TMEM35A significantly reduced the GFP-positive fraction by approximately 70% and 50%, respectively, consistent with impaired cellular fitness (Fig. 2g,h). In contrast, *FBXW11* targeted cells were enriched within organoids by approximately 34%, supporting a cell competitive advantage following *FBXW11* loss.

Together, our validation results confirmed that ORIGAMI can detect gene regulators for cerebral organoid development. Of these, given the robust and reproducible organoid overgrowth phenotype associated with *FBXW11* loss, we next sought to define its role in regulating human brain development.

### *FBXW11* deficiency leads to a macrocephalic phenotype

*FBXW11* encodes an F-box/WD-repeat protein that functions as the substrate recognition component of the SCF ubiquitin ligase complex. To determine whether organoid overgrowth phenotype was specific to *FBXW11* disruption rather than a guide-specific effect, we generated a second independent sgRNA targeting a distinct exonic region. The original and second sgRNA produced 69% and 75% editing efficiencies, as determined by Sanger sequencing trace analysis (Fig. 3a). When differentiated into cerebral organoids, both *FBXW11*-targeted iPSCs populations reproduced enlarged organoids compared to controls (Fig. 3b,c), supporting that our observations are not the results of a specific off-target effect from the original guide. By quantifying the area occupied by individual cerebral organoids, we were able to detect larger organoids as early as day 5 (for sgRNA2), and on days 10, 20, and 40 for both guides (Fig. 3c). In wild-type organoids, *FBXW11* mRNA expression gradually increased across differentiation (Fig. 3d), mirroring the decline in proliferation observed at later stages of neurodevelopment. Supporting this, *FBXW11* overexpression reduced embryoid body/organoid size at early stages (Fig. 3e-g), the opposite of the knockout phenotype. Together, these gain- and loss-of-function data support a role for *FBXW11* as a negative regulator of cerebral organoid growth.

**Fig. 3.**
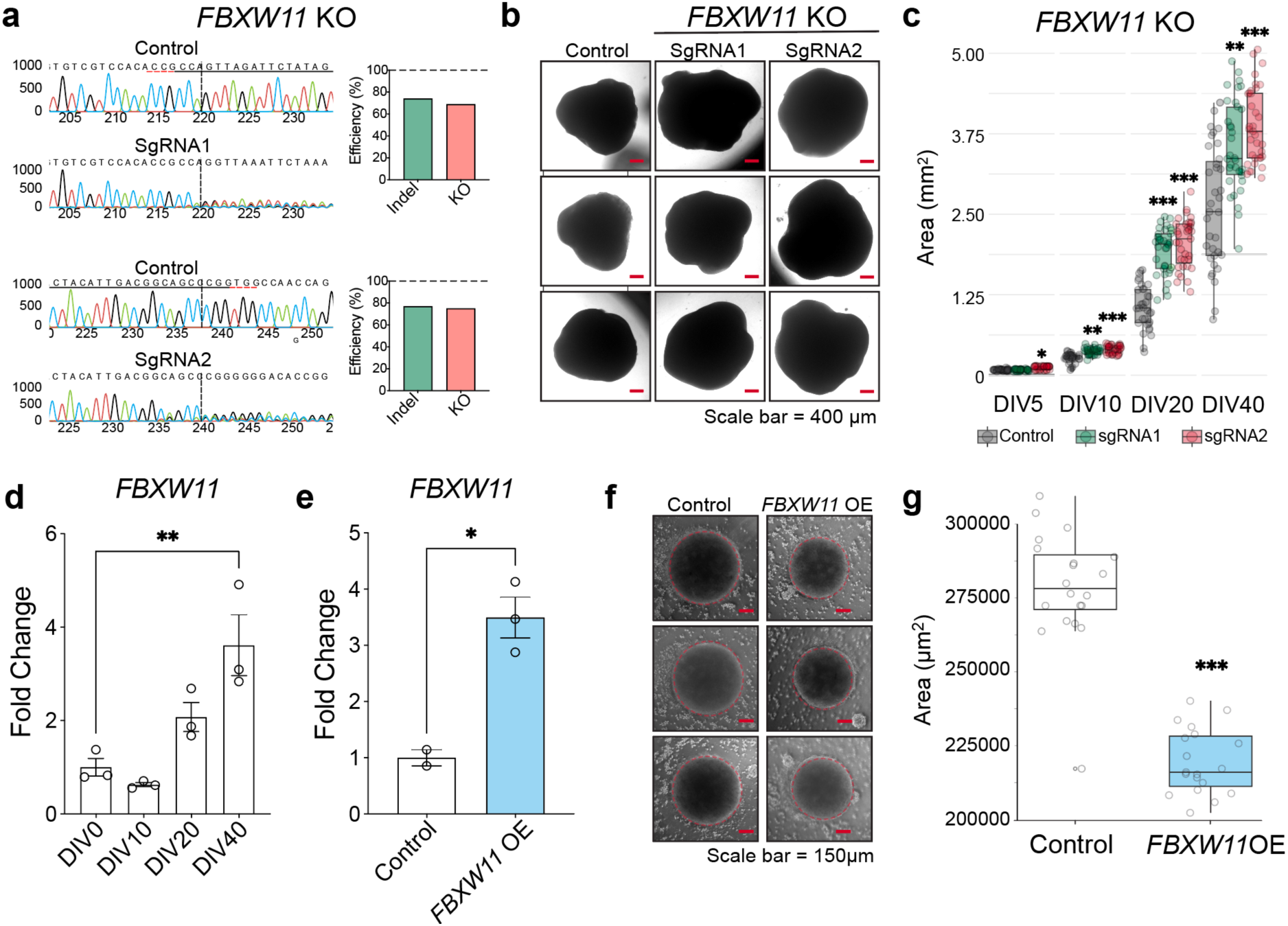
*FBXW11* restricts cerebral organoid growth during early neurodevelopment. **a,** Validation of CRISPR-mediated *FBXW11* knockout in human pluripotent stem cells. Representative Sanger sequencing traces, and ICE-based editing efficiencies for independent sgRNAs relative to controls. **b,** Representative brightfield images of cerebral organoids (DIV40) generated from control and *FBXW11* KO iPSC populations across independent differentiations. *FBXW11* deficient organoids display increased size relative to controls. **c,** Quantification of organoid area during differentiation at DIV5, DIV10, DIV20, and DIV40. Each dot represents an individual organoid pooled from n=3 independent differentiations. Boxplots indicate median and interquartile range. Statistical significance was determined by two-way ANOVA; *P < 0.05, **P < 0.01, ***P < 0.001. **d,** *FBXW11* expression measured by RT-qPCR during cerebral organoid differentiation. Each point represents pooled organoids from an independent differentiation. Data are presented as fold change relative to DIV0. N=3 independent differentiations. Statistical significance was determined by one-way ANOVA; **P < 0.01. **e,** RT-qPCR validation of *FBXW11* expression in overexpression engineered iPSCs compared with control cells. Statistical significance was determined by unpaired t test; *P < 0.05. **f,** Representative brightfield images of embryoid bodies/organoids generated from control and FBXW11 overexpressing iPSCs. **g,** Quantification of DIV5 embryoid bodies/organoids size following *FBXW11* overexpression. Each dot represents an individual organoid pooled from n=3 independent differentiations. Statistical significance was determined by two-way ANOVA; *P < 0.05, **P < 0.01, ***P < 0.001.

We proceeded to characterize *FBXW11* deficiency, specifically we asked whether the organoid overgrowth reflected proliferation already present in the stem cell stage. Cell counts across multiple culture days showed no evidence of increased proliferation in *FBXW11* deficient iPSCs (Extended Data Fig. 2a), although *FBXW11* overexpression modestly reduced stem-cell expansion (Extended Data Fig. 2b). BrdU pulse-labelling followed by flow cytometry revealed comparable proportions of cells in G1, S, and G2/M phases between *FBXW11* KO and control iPSCs (Extended Data Fig. 2c,d), further indicating that *FBXW11* loss does not strongly alter cell-cycle regulation at the pluripotent stage. RNA-seq of *FBXW11* KO iPSCs identified 392 downregulated and 323 upregulated genes at statistical significance (adjusted P < 0.05), although most changes were modest with only 21 genes showing an absolute fold change greater than two (Extended Data Fig. 2e; Extended Data Table 2). Reduced *FBXW11* transcript abundance was among the strongest differentially expressed signals, consistent with effective CRISPR targeting. Top selected differentially expressed genes were validated by RT-qPCR across both sgRNA guides (Extended Data Fig. 2f). Importantly, this transcriptomic dataset did not reveal a broad activation of proliferation programs that could explain the cerebral organoid phenotype. These findings suggest that *FBXW11* does not primarily regulate pluripotent stem-cell cell division but instead acts during neurodevelopmental differentiation to restrict organoid expansion.

### Single-cell RNA-Seq reveals expansion of radial glia and impaired neuronal maturation in *FBXW11*-deficient cerebral organoids

Because *FBXW11* loss produced only modest changes in pluripotent stem cells, we next focused on DIV40 cerebral organoids, where the overgrowth phenotype was most pronounced (Fig. 3c). To characterize the consequences of complete *FBXW11* deficiency, we isolated iPSC knockout clones. These lines showed single base insertions predicted to cause a frameshift with robust KO scores (99%), and they recapitulated the organoid size increase observed in pooled CRISPR-edited cultures (Extended Data Fig. 3a,b). Using these clones we made cerebral organoids and subjected them to single-cell RNA-seq (see Methods). After demultiplexing and quality filtering (removing doublets, cells with >15% mitochondrial reads, and cells outside defined gene and UMI thresholds), the two pools were integrated with Harmony (Extended Data Fig. 3c-g).

To resolve the cellular composition of the organoids, we projected the integrated single-cell data onto a UMAP, a two-dimensional embedding in which each point is a single cell and cells with similar transcriptomes cluster together (Fig. 4a,b). Unbiased clustering (Leiden, resolution 0.4) resolved distinct neurodevelopmental populations, which we annotated by reference-based label transfer from a published human cerebral organoid atlas^16^ (see Methods) as radial glia, neural progenitor cells (NPCs), neurons, immature choroid plexus/hem populations, and mature choroid plexus-like cells (Fig. 4a,b). When the UMAP colour by genotype, control and *FBXW11*-KO cells occupy overlapping regions, despite the loss of *FBXW11*, indicating preservation of overall developmental identity (Fig. 4c). However, the proportion of cells in each quantification shifted significantly (Pearson’s Chi-squared test, X^2^=869.57, df=6, p<2.2×10^-16^). Quantifying cell-type composition using standardised Pearson residuals revealed a marked expansion of radial glia populations in FBXW11 KO organoids (Fig 4d, Extended Data Table 3, residual = 19.08), accompanied by relative reductions in differentiated neuronal populations and choroid plexus-associated lineages (Fig. 4d, all ChP residuals < -2.9). Correlation analysis of highly variable genes (n = 2,000) confirmed cell-types clustered by type, not genotype, while highlighting condition-specific transcriptional shifts were most distinct within radial glia and neuronal compartments (Fig. 4e). These data suggest that *FBXW11* deficiency selectively remodels the balance between progenitor and neuronal states within developing cerebral organoids.

**Fig. 4.**
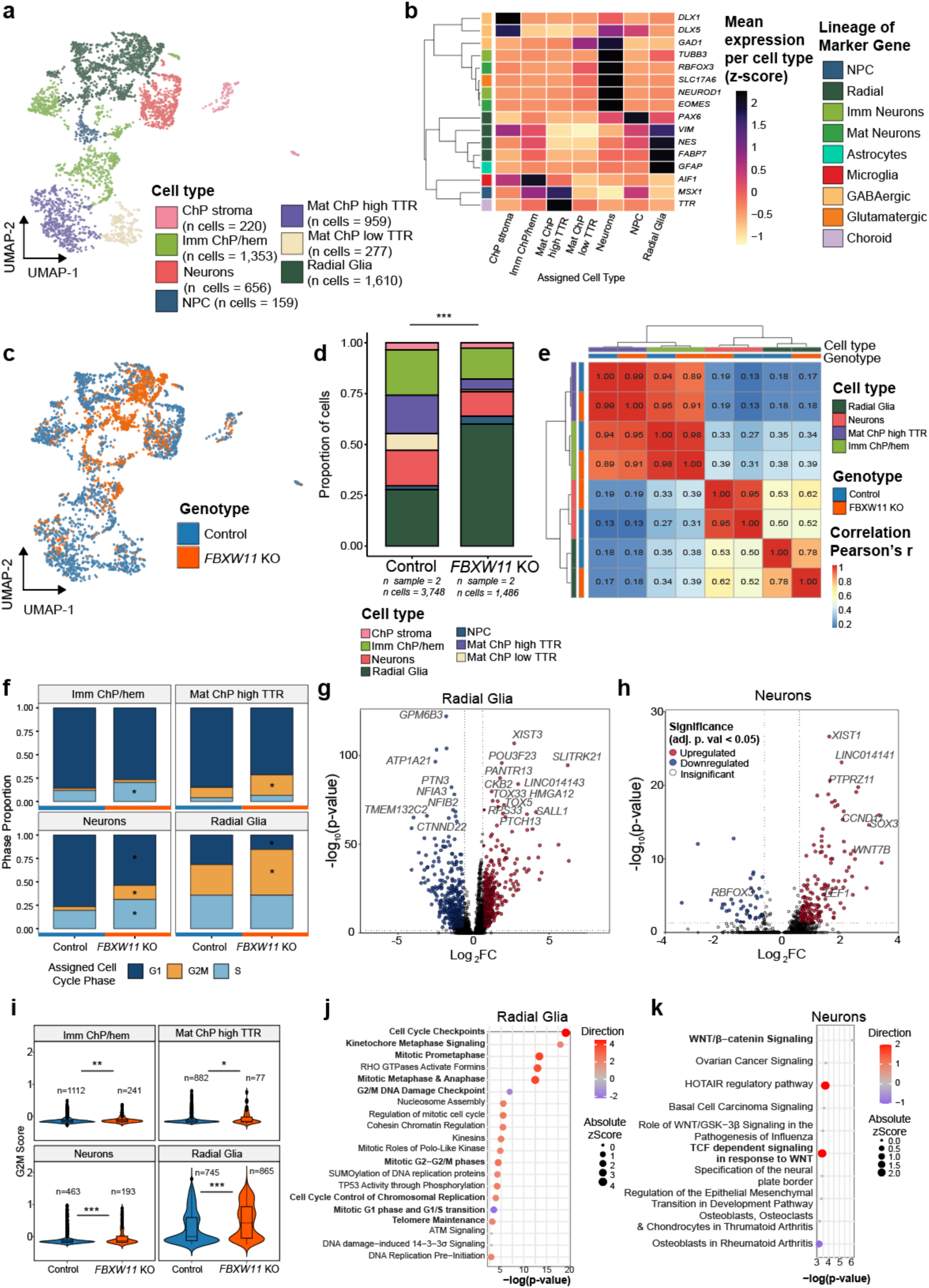
Single-cell RNA-Seq in of KO organoids *FBXW11* KO organoids reveals altered cell-state composition, proliferative programs, and dysregulated neuronal WNT/β-signalling. **a,** UMAP (Uniform Manifold Approximation and Projection) of integrated single-cell RNA-seq data showing major cell populations identified across all samples. Clusters were annotated by reference-based label transfer into choroid plexus stroma (ChP stroma), immature choroid plexus/hem (Imm ChP/hem), mature choroid plexus high transthyretin-expressing cells (Mat ChP high TTR), neurons, neural progenitor cells (NPCs), and radial glia. **b,** Heatmap of representative marker genes used for cluster annotation. Colour indicates mean expression per cluster z-scored across clusters for each gene (magma scale; yellow = low, purple/black = high, range approximately −1.5 to +2.5). Genes are hierarchically clustered by expression profile (row dendrogram); the adjacent annotation bar groups the same genes by gene marker lineage. Cell types are shown on the x-axis. Markers shown are *PAX6*, *VIM*, *NES* and *FABP7* (radial glia), *MSX1* (neural progenitor), *EOMES* and *NEUROD1* (neurogenic), *TUBB3* (immature neuron), *RBFOX3* (mature neuron), *SLC17A6* (glutamatergic), *GAD1*, *DLX1* and *DLX5* (GABAergic), *GFAP* (astroglia), *AIF1* (microglia), and *TTR* (choroid plexus). **c,** The same UMAP as Fig. 4a, coloured by genotype (Control, blue; *FBXW11* KO, green), showing that both genotypes occupy overlapping regions of transcriptional space. **d,** Stacked bar plot showing the proportion of cells assigned to each cell type in Control and FBXW11 KO organoids. *FBXW11* KO organoids display an increased proportion of radial glia and altered neuronal/choroid plexus populations relative to controls. Differences in overall cell type distributions were evaluated using a Pearson’s Chi-squared test on pooled sample counts, with specific population shifts identified by standardized Pearson residuals > |2|. Data obtained from *n* = 3 independent differentiations (see methods) comprising 3,748 Control cells and 1,486 *FBXW11* KO cells. Abbreviations: ChP, choroid plexus; NPC, neural progenitor cell; TTR, transthyretin. **e,** Correlation heatmap of highly variable genes across cell types and sample conditions. Hierarchical clustering demonstrates stronger transcriptional similarity within cell types than between experimental conditions, while revealing condition-specific shifts in radial glia and neuronal populations. **f,** Cell-cycle phase distribution across major cell populations in Control and *FBXW11* KO organoids. Volcano plots showing differentially expressed genes (DEGs) in radial glia (**g**) and neurons (**h**) comparing *FBXW11* KO versus Control organoids. Differential expression was performed within each cell type using Seurat’s FindMarkers (Wilcoxon rank-sum test implemented via limma, wilcox_limma; min.pct = 0.25) with Benjamini–Hochberg adjustment. Axes show log2 fold-change (x) versus −log10 adjusted p-value (y); points are coloured as upregulated (skyblue) or downregulated (pink) at adjusted p < 0.05, or not significant (grey). **i,** Violin with overlapping box plots of per-cell G2/M cell-cycle scores across cell types with >500 cells, comparing Control and *FBXW11* KO. Genotypes were compared by two-sided Wilcoxon rank-sum test (p-values and per-group n shown on each panel); G2/M scores were significantly elevated in KO radial glia (p < 2.2 × 10⁻¹⁶) and neurons. Pathway enrichment analysis of radial glia (**j**) and neuron (**k**) DEGs, respectively.

Given the expansion of progenitor-like populations, we next examined proliferative signatures across annotated cell types directly. *FBXW11* KO organoids showed a marked depletion of G1-phase cells and a corresponding increased proportions of G2/M-phase cells within radial glia, maturing choroid plexus and neuronal populations (Fig. 4f, cell types with >500 total cells). G2/M scores were correspondingly and significantly elevated, consistent with enhanced progenitor proliferation or delayed cell-cycle exit. Pseudobulk differential expression analysis of radial glia pseudobulk population revealed strong upregulation of genes associated with proliferation and neurodevelopmental programs, whereas neuronal populations showed altered expression of differentiation-associated transcripts (Fig. 4g). Consistent with these findings, pathway enrichment analysis revealed significant activation of mitotic, DNA replication, and checkpoint pathways in radial glia from *FBXW11* KO organoids (Fig. 4j), while neuronal populations displayed altered WNT/β-catenin and developmental signalling pathways (Fig. 4h,k). Notably, this included reduced expression of *RBFOX3* (encoding NeuN), a canonical marker of mature, post-mitotic neurons, consistent with a shift from terminal neuronal differentiation. Collectively, this data suggests that loss of *FBXW11* promotes expansion and proliferative maintenance of radial glia-like progenitors at the expense of neuronal maturation, providing a potential mechanistic explanation for the enlarged organoid phenotype.

### FBXW11 regulates cortical maturation and cytoarchitecture

To validate the transcriptional changes identified by single-cell RNA-seq, we next examined progenitor and neuronal markers in DIV40 cerebral organoids by immunofluorescence. Consistent with the expanded radial glia population observed in the single-cell dataset, *FBXW11* KO organoids displayed enlarged SOX2- and PAX6-positive neuroepithelial domains compared with controls (Fig. 5a, Extended Data Fig. 4a). Quantification of SOX2-positive nuclei confirmed a significant increase across both independent *FBXW11* KO lines relative to control organoids (Fig. 5b), supporting the conclusion that *FBXW11* deficiency promotes maintenance or expansion of neural progenitor-like populations.

**Fig. 5.**
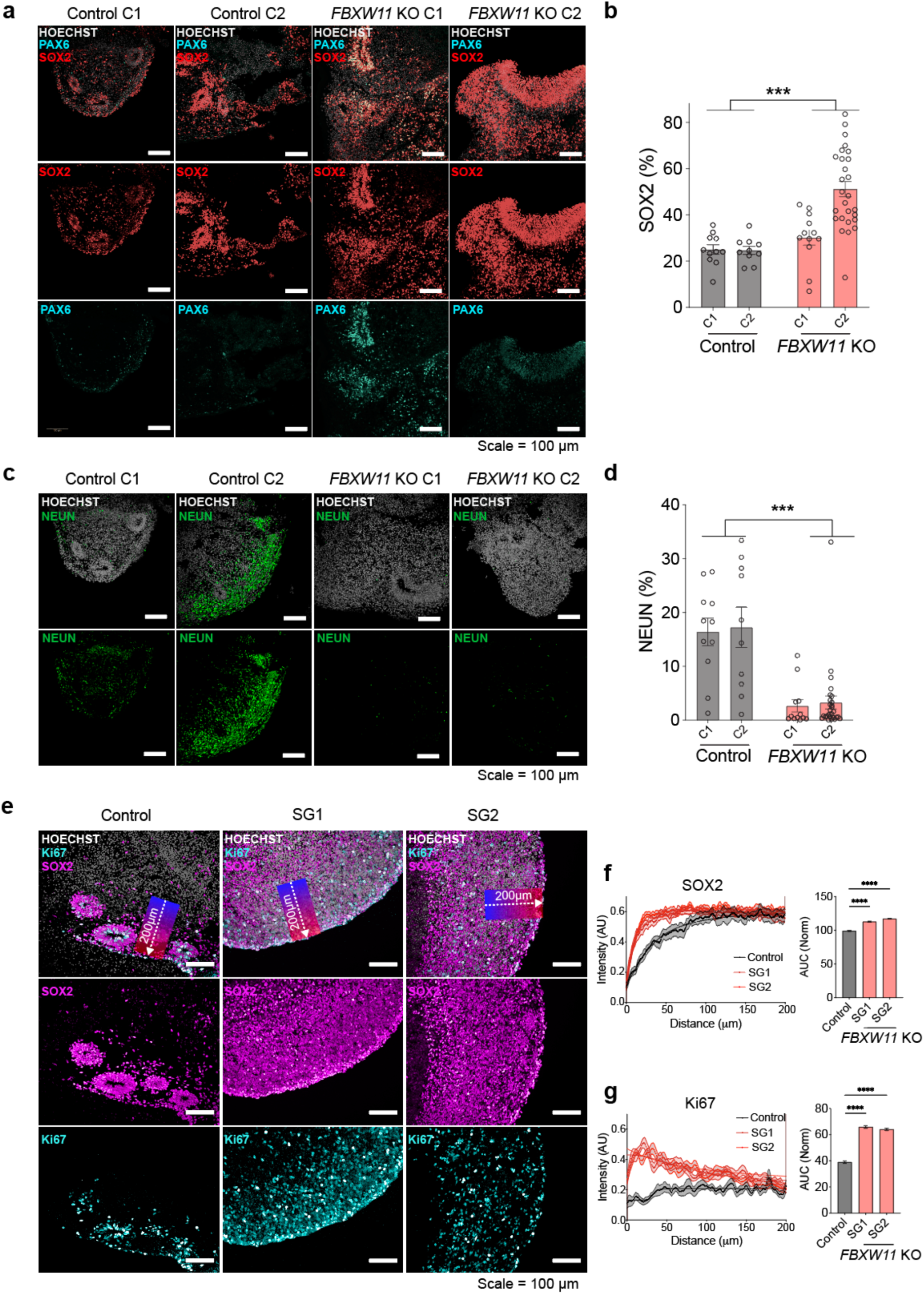
FBXW11 deficiency promotes expansion of SOX2/PAX6-positive progenitor domains and loss of neuronal differentiation in DIV40 cerebral organoids. **a,** Representative immunofluorescence images of DIV40 Control and *FBXW11* KO cerebral organoids stained for the neural progenitor markers SOX2 and PAX6. *FBXW11* KO organoids display expanded SOX2- and PAX6-positive neuroepithelial regions relative to controls. **b,** Quantification of SOX2-positive nuclei across Control and *FBXW11* KO organoids. Each dot represents an individual organoid across n = 3 independent differentiations. Statistical significance was determined by two-way ANOVA; **P < 0.001. **c,** Representative immunofluorescence images of DIV40 organoids stained for the neuronal marker NeuN. **d,** Quantification of NeuN-positive nuclei in Control and *FBXW11* KO organoids. Each dot represents an individual organoid across n=3 independent differentiations. Statistical significance was determined by two-way ANOVA; **P < 0.001. **e,** Representative immunofluorescence images of DIV40 organoids stained for SOX2 and the proliferation marker Ki67. Insets illustrate representative radial intensity measured regions extending 200 μm from the ventricular edge toward the outer organoid layer. **f,** Radial intensity profiling of SOX2 across ventricular-like regions in Control and *FBXW11* KO organoids, and Area-under-the-curve (AUC) quantification. Statistical significance was determined by one-way ANOVA; ****P < 0.0001. **g,** Radial intensity profiling of Ki67 across ventricular-like regions in Control and *FBXW11* KO organoids, and AUC quantification. Statistical significance was determined by one-way ANOVA; ****P < 0.0001.

In contrast, staining for the neuronal maturation marker NeuN revealed a marked reduction in differentiated neuronal populations within *FBXW11* KO organoids (Fig. 5c). Quantification of NeuN-positive nuclei showed a significant decrease in neuronal across both KO clones relative to controls (Fig. 5d), consistent with the reduced neuronal populations identified by single-cell RNA-seq and with decreased RBFOX3 expression in *FBXW11*-deficient neurons (Fig. 4h). We also observed reduced expression of the cortical neuronal marker CTIP2 in KO organoids (Extended Data Fig. 4b). Together these findings indicate that loss of *FBXW11* impairs neuronal maturation while maintaining organoids in a more progenitor-like state.

Given the increased progenitor abundance and elevated cell-cycle signatures identified in the scRNA-seq analysis, we next examined proliferative organization within ventricular-like regions of the organoids. Co-staining for SOX2 and Ki67 demonstrated pronounced expansion of proliferative neuroepithelial zones in *FBXW11* KO organoids relative to controls (Fig. 5e). Radial intensity profiling extending 200 μm from the ventricular edge toward the outer organoid layer consistently found elevated SOX2 signal throughout the neuroepithelial compartment in knockouts (Fig. 5f). Similarly, Ki67 intensity profiles coincided with a broad increase in proliferation across the expanded ventricular-like regions of *FBXW11* KO organoids (Fig. 5g). Area-under-the-curve (AUC) quantification of these two markers statistically supported an increase in proliferation and glial/progenitor markers. Collectively, these data support a model in which *FBXW11* loss promotes expansion and proliferation of radial glia-like progenitors while suppressing neuronal differentiation, appropriate cortical layer formation, and ultimately contributing to the enlarged organoid phenotypes.

### FBXW11 regulates ventricular rosette organization during cerebral organoid development

After observing expanded progenitor domains and altered proliferative organization in DIV40 *FBXW11* KO organoids, we next asked whether *FBXW11* loss affected the formation and maturation of ventricular-like rosettes during early stages of cerebral organoid development. Rosette structures are hallmark neuroepithelial features of cerebral organoids and model key aspects of ventricular zone organization during cortical development^17^. To assess rosette architecture, we stained DIV10 and DIV20 organoids for SOX2, β-catenin (CTNNB1), and TUBB3 to visualize progenitor domains, apical junctional organization, and early neuronal differentiation, respectively. At DIV10, *FBXW11* deficient organoids displayed an increased number of ventricular-like rosettes relative to controls (Fig. 6a,c). However, rosette density normalized to organoid area was not significantly altered (Fig. 6d), indicating that the increased rosette number was largely proportional to the increased size of KO organoids. At this time point, TUBB3 intensity was also reduced in FBXW11 KO organoids (Extended Data Fig. 4c,d), supporting the conclusion that early neuronal differentiation programs are already disrupted before the pronounced neuronal maturation defects observed at DIV40 (Fig. 5d, Extended Data Fig. 4).

**Fig. 6.**
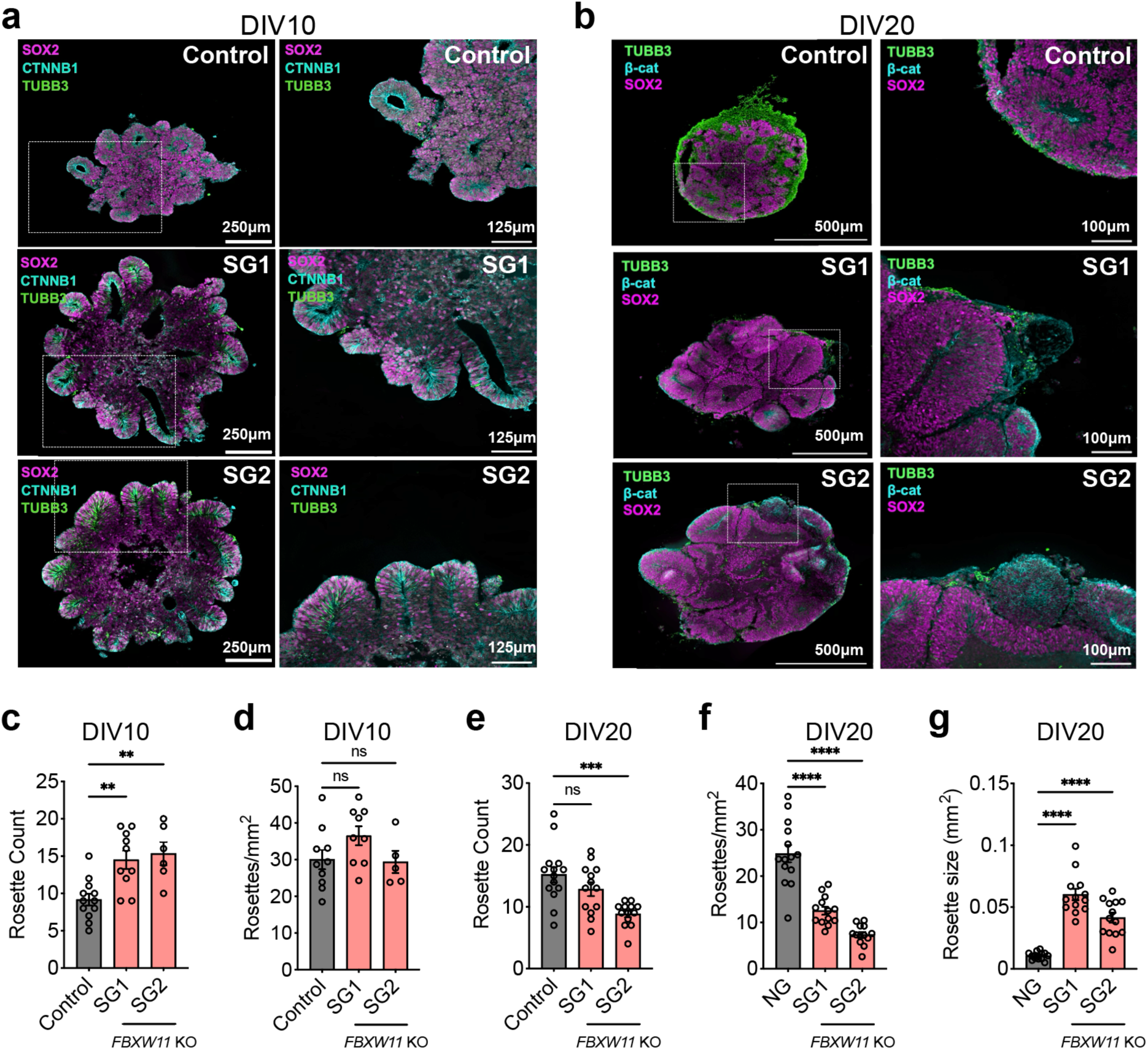
*FBXW11* deficiency alters ventricular rosette organization during early cerebral organoid development. **a,** Representative immunofluorescence images of DIV10 cerebral organoids stained for SOX2, CTNNB1 (β-catenin), and TUBB3. **b,** Representative immunofluorescence images of DIV20 cerebral organoids stained for SOX2, β-catenin, and TUBB3. **c,** Quantification of ventricular-like rosette number per organoid at DIV10. Each dot represents an individual organoid pooled across 3 experimental replicates. Statistical significance was obtained using one-way ANOVA; **P < 0.001. **d,** Quantification of rosette density (rosettes/mm²) at DIV10. Each dot represents an individual organoid pooled across 3 experimental replicates. **e,** Quantification of ventricular-like rosette number per organoid at DIV20. Each dot represents an individual organoid pooled across 3 experimental replicates. Statistical significance was obtained using one-way ANOVA; **P < 0.001.**f,** Quantification of rosette density (rosettes/mm²) at DIV20. Each dot represents an individual organoid pooled across 3 experimental replicates. Statistical significance was obtained using one-way ANOVA; ****P < 0.0001.**g,** Quantification of average rosette size at DIV20. Each dot represents an individual organoid pooled across 3 experimental replicates. Statistical significance was obtained using one-way ANOVA; ****P < 0.0001.

By DIV20, structural differences between control and *FBXW11* KO organoids became more pronounced (Fig. 6b). Total rosette number was reduced in *FBXW11* SG2 organoids and trended lower in *FBXW11* SG1 organoids (Fig. 6e), while rosette density was significantly reduced in both KO lines (Fig. 6f). This reduction in density was accompanied by a marked increase in average rosette size (Fig. 6g), consistent with expansion and possible merging of ventricular-like neuroepithelial domains over time. High-magnification imaging further showed enlarged SOX2-positive regions organized around β-catenin-enriched apical structures in *FBXW11* KO organoids (Fig. 6b).

Together, these results indicate that *FBXW11* regulates the spatial organization and maturation of ventricular-like rosettes during cerebral organoid development. Loss of *FBXW11* promotes early expansion of neuroepithelial structures, followed by the formation of enlarged progenitor-rich ventricular domains, consistent with the increased proliferative radial glia state identified by single-cell transcriptomic and immunohistochemistry.

### FBXW11 interacts with β-catenin and negatively regulates WNT signalling during cerebral organoid development

To identify mechanisms by which *FBXW11* regulates neurodevelopmental programs, we engineered an FBXW11-TurboID fusion construct and expressed it in stem cells to label proteins in close proximity to FBXW11 (Fig. 7A). Biotinylated proteins were isolated and identified by mass spectrometry and compared to control samples (Fig. 7b,c). FBXW11 peptides were not detected in control lysates, but were enriched in FBXW11-TurboID expressing fractions (Extended Data Fig. 4e), supporting that our isolation protocol and mass spectrometry enriched for our target protein. β-catenin, encoded by *CTNNB1*, was the top identified FBXW11 proximal protein, supporting a connection between FBXW11 and canonical WNT signalling. To validate this interaction independently, we performed co-immunoprecipitation of HA-tagged FBXW11, which confirmed enrichment of FBXW11 in HA immunoprecipitates relative to IgG controls and demonstrated co-enrichment of β-catenin (Fig. 7c,d).

**Fig. 7.**
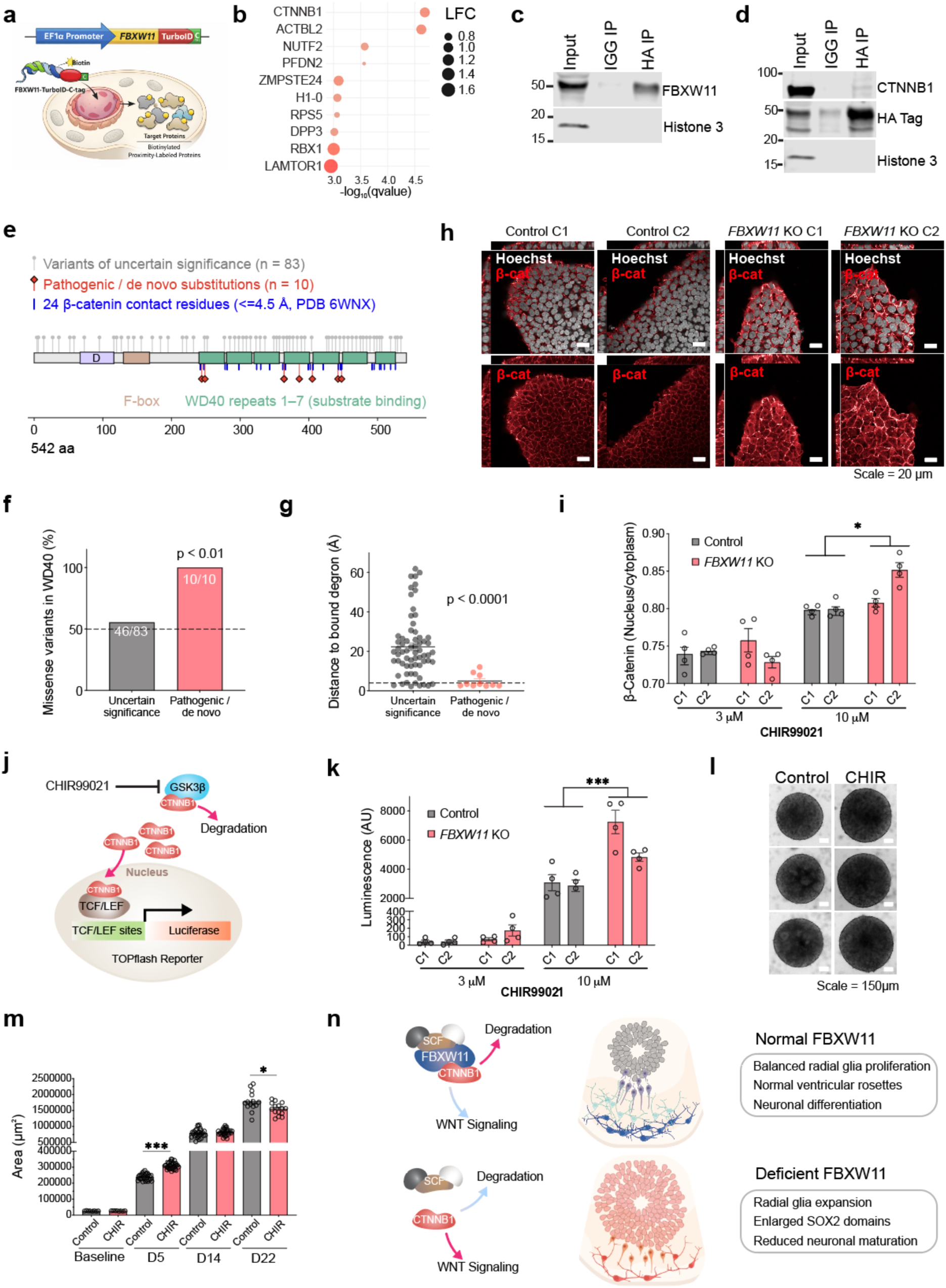
FBXW11 interacts with β-catenin and negatively regulates WNT signalling during cerebral organoid development. **a,** Schematic showing FBXW11-TurboID proximity labelling strategy. **b,** Top enhanced proteins identified by FBXW11-TurboID proximity labelling compared to controls. **c,** Validation of HA-tagged FBXW11 immunoprecipitation. Immunoblotting confirms efficient enrichment of FBXW11 in HA immunoprecipitated fraction. **d,** Co-immunoprecipitation analysis of FBXW11 and β-catenin. Cells expressing HA-tagged FBXW11 were subjected to HA immunoprecipitation. Immunoblotting for CTNNB1 demonstrates association of β-catenin with FBXW11. **e,** Domain architecture of FBXW11 (UniProt Q9UKB1) with all mappable missense variant plotted at their residues. Vertical ticks in the WD40 band mark the 24 residues within 4.5 Angstrom of the bound β-catenin degron in PDB 6WNX (24 residues). **f,** Proportion of missense variants within the WD40 repeats by class. Dashed line = expected proportion (49.3 percent, 267 of 542 residues). Statistical significance determined by Fisher exact one-sided test. **g,** Minimum heavy-atom distance from each variant residue to the bound degron peptide in PDB 6WNX. Dotted line = 4 Å direct-contact threshold fixed a priori. Statistical significance determined by Mann-Whitney one-sided. **h,** Representative immunofluorescence images of Control and FBXW11 KO cells treated with the GSK3β inhibitor CHIR99021 (10 μM). **i,** Quantification of β-catenin nuclear localisation following treatment with CHIR99021 (3 μM or 10 μM). Data are expressed as the nuclear-to-cytoplasmic β-catenin intensity ratio. n = 4 independent experiments. Bars represent mean ± SEM. Statistical significance was determined by two-way ANOVA (*, P < 0.05). **j,** TOPFlash luciferase reporter assay^19^ measuring canonical WNT/β-catenin signalling activity following CHIR99021 treatment, an inhibitor of GSK3β allowing accumulation of CTNNB1 and increased luciferase reporter signal. **k,** Luminescence measured in clonal control and *FBXW11* KO iPSCs. Data are presented as mean ± SEM. Statistical significance was determined by two-way ANOVA; **P < 0.001. **l,** Representative brightfield images of cerebral organoids cultured in the presence or absence of CHIR99021 (10 μM). **m,** Quantification of organoid area during differentiation in untreated and CHIR99021-treated conditions. Each dot represents an individual organoid. Bars represent mean ± SEM. Statistical significance was determined by two-way ANOVA; *P < 0.05, **P < 0.001. **n,** Schematic showing proposed mechanism for FBXW11 function in cerebral organoids.

Mapping variants of uncertain significance and pathogenic/de novo substitutions onto the FBXW11 protein structure showed that pathogenic/de novo substitutions were concentrated within the WD40 repeat region, which contains the predicted β-catenin substrate-binding interface (Fig. 7e,f), an observation first reported by ^18^. Since then, an experimental structure of FBXW11 bound to its substrate has since become available (PDB 6WNX). Using this we found that pathogenic/de novo substitutions were significantly closer to β-catenin contact residues (median 4.7Å from the doubly phosphorylated β-catenin degron) than variants of uncertain significance (Fig, 7g). These observations support the clinical relevance of the *FBXW11* substrate-binding domain and suggest that disruption of β-catenin recognition may contribute to developmental phenotypes associated with *FBXW11* variation.

We therefore examined β-catenin localization in *FBXW11*-deficient cells. Under basal conditions, we did not detect clear differences in β-catenin abundance or localization (Data not shown). However, treatment with the GSK3β inhibitor CHIR99021 induced β-catenin nuclear translocation in a dose-dependent manner, and this response was enhanced in *FBXW11* KO cells relative to controls (Fig. 7h,i). These data suggest that FBXW11 deficiency sensitizes cells to WNT pathway activation rather than producing a large constitutive increase in β-catenin under baseline conditions.

To test whether this altered β-catenin localization translated into increased pathway activity, we transfected cells with the TOPFlash WNT/β-catenin reporter^19^ and measured luciferase activity (Fig. 7j). Basal reporter activity was below the threshold of detection, whereas CHIR99021 treatment increased TOPFlash activity in a dose-dependent manner (Fig. 7j,k). This induction was significantly enhanced in *FBXW11*-deficient cells, indicating that FBXW11 restrains β-catenin-dependent transcriptional responses under conditions of WNT pathway stimulation.

We next asked whether pharmacological WNT activation could mimic the organoid overgrowth phenotype observed following *FBXW11* loss. Moderate CHIR99021 treatment increased cerebral organoid size at early stages of differentiation, including DIV5 (Fig. 7l,m), although this effect was not sustained at later time points. Collectively, these data support a model in which FBXW11 physically associates with β-catenin and constrains WNT/β-catenin signalling, thereby limiting radial glial expansion and cerebral organoid growth (Fig. 7n).

## Discussion

In this study we aimed to identify underexplored targets with probable function in the human brain. To do this, we took advantage of comparative genomics by collecting genes causing lethality when disrupted in the fly’s nervous system. These were further filtered to enrich by reduced knowledge, which we defined by limited gene ontology annotations. High confidence human orthologs were then engineered into a CRISPR library encompassing 129 experimental genes, which were tested using cerebral organoids throughout early stages of development.

Among these candidates, *FBXW11* emerged as a robust regulator of cerebral organoid development. Loss of *FBXW11* increased organoid size, expanded radial glia populations, enhanced cell-cycle signatures, reduced neuronal maturation, and altered ventricular rosette architecture. These phenotypes converge on a central developmental principle: cortical growth depends on the balance between neural progenitor self-renewal and neurogenic commitment. Human-specific features such as basal radial glia, ARHGAP11B, NOTCH2NL, and human accelerated regulatory elements are thought to contribute to neocortical expansion by shifting this balance ^20-23^. The same axis is disrupted in disease, where gain-of-function mutations in the PI3K-AKT-mTOR pathway can cause megalencephaly and hemimegalencephaly, whereas loss-of-function CTNNB1 variants are associated with microcephaly and neurodevelopmental impairment^24,25^. Our findings place FBXW11 within this broader framework as a negative regulator of progenitor expansion and tissue growth during early human neurodevelopment.

Mechanistically, our data support a model in which FBXW11 constrains WNT/β-catenin signalling during cerebral organoid development. β-catenin is a dose-sensitive regulator of cortical size: stabilizing β-catenin in neural precursors enlarges the mouse brain, and persistent β-catenin activity expands SOX2+/PAX6+ progenitors at the expense of neuronal differentiation ^26,27^. Consistent with this biology, FBXW11-TurboID proximity labelling identified CTNNB1 as a high-confidence FBXW11-associated protein, FBXW11-deficient cells showed enhanced β-catenin nuclear localization and TOPFlash reporter activation after WNT stimulation, and pharmacological WNT activation partially reproduced the early organoid overgrowth phenotype. Together, these observations support that while FBXW11 does not affect proliferation at the pluripotent stem cell stage, it helps tune developmental signalling during the transition into organized neuroepithelial tissue.

Clinically, rare *de novo* variants in FBXW11 have been reported in individuals with neurodevelopmental disorders, including developmental delay, intellectual disability, autism spectrum disorder, epilepsy, and craniofacial abnormalities (OMIM #618914)^18,28^. Although the clinical spectrum remains incompletely defined and brain overgrowth has not emerged as a consistent phenotype, these observations support an essential role for FBXW11 during human development. Our organoid data provide a potential cell-biological framework for understanding these disorders by demonstrating that loss of FBXW11 disrupts the balance between progenitor expansion and neuronal differentiation through enhanced WNT/β-catenin responsiveness. Given the dosage sensitivity of WNT/β-catenin signalling during corticogenesis, even partial changes in FBXW11 activity may alter developmental trajectories in ways that contribute to patient phenotypes. Future studies using patient-derived iPSCs and variant-specific knock-in organoids will be important for determining how individual FBXW11 mutations affect protein function, pathway regulation, and cortical development.

Cerebral organoids derived from human induced pluripotent stem cells (iPSCs) provide an experimentally tractable system for forward genetic dissection of human cortical development^17^. However, pooled perturbation screens in three-dimensional tissues face important technical limitations: mosaic transduction, variable organoid growth, and stochastic clonal expansion. which can generate apparent sgRNA enrichment or depletion that does not reflect a reproducible biological phenotype. CRISPR-LICHT addressed this challenge by coupling sgRNAs to lineage barcodes and using PCR-derived cell barcodes to estimate lineage size at cellular resolution in organoids^8^. Subsequent approaches have extended pooled perturbation screening to single-cell organoid assays and assembloid systems ^29-31^. Here, we developed ORIGAMI as a complementary and experimentally simple strategy based on a modified LentiCRISPRv2 architecture^32^. Rather than reconstructing lineage size at single-cell resolution, similar to CRISPR-LICHT, ORIGAMI uses barcode diversity as internal replication for each sgRNA. True biological hits are expected to be supported by many independently represented barcodes, whereas apparent enrichment driven by only a small number of barcoded clones can be flagged as a potential founder effect. Because sgRNA and barcode abundance are recovered using standard pooled CRISPR library prep followed by short-read sequencing, ORIGAMI provides a low-barrier approach for pooled genetic screening in heterogeneous three-dimensional models.

Previous organoid and assembloid CRISPR screens have successfully interrogated genes nominated from ASD exomes^29^, microcephaly genetics^8^, or GWAS loci ^33^. These approaches have been highly informative, but they naturally focus on genes already connected to human disease. In contrast, we curated candidates from *Drosophila* nervous-system lethality screens and prioritized conserved genes with limited functional annotation, reasoning that this strategy could expose underexplored regulators of human neurodevelopment. Our results support this rationale. Although *Drosophila* and mammalian brains differ substantially in scale and architecture, many pathways governing neural development are deeply conserved. By combining comparative genetics with cerebral organoid screening, we identified both expected regulators and previously understudied candidates with strong effects on human organoid growth.

Several limitations of this study should be considered. Cerebral organoids incompletely recapitulate later developmental stages and lack aspects of vascularization and environmental inputs present in vivo. Furthermore, although our data support β-catenin as a major downstream effector of *FBXW11*, additional substrates identified through proximity proteomics may also contribute to the observed phenotype. Future studies examining these pathways in more mature organoid systems and *in vivo* models will help further define the developmental functions of *FBXW11*.

In summary, we establish *FBXW11* as a regulator of human neurodevelopment that restricts progenitor expansion and promotes neuronal maturation through modulation of WNT/β-catenin signalling. More broadly, our work demonstrates how comparative genomics coupled with barcode-enabled pooled CRISPR screening in human organoids can accelerate the discovery of conserved regulators of brain development.

## Methods

### Screening target curation

Genome-wide *Drosophila* nervous-system RNAi lethality datasets were used to identify genes required for central nervous system development ^4,5^. Drosophila genes whose knockdown produced lethality using either the pan-neuronal elav-Gal4 driver or the neural stem cell inscuteable-Gal4 driver were curated through two convergent prioritization pipelines. In the first pipeline, *Drosophila* genes were filtered for unknown or unannotated FlyBase Gene Ontology assignments^9^, and retained if they had high-confidence human orthologues defined by DIOPT score >10^10^. In the second pipeline, high-confidence human orthologues were first identified and then filtered for limited human Gene Ontology annotation using GO::TermFinder ^11^. Candidate genes from both pipelines were merged to generate the final screening library. Additional positive and negative control genes were included, including genes with established roles in pluripotency, proliferation, or neurodevelopment. The complete gene list and sgRNA sequences are provided in Extended Data Table 1.

### Library preparation

For ORIGAMI screening, LentiCRISPRv2 (Addgene #52961) was modified to include a random 5-nucleotide barcode downstream of the sgRNA cassette. The parental plasmid was digested with EcoRI and NheI, purified, and circularized using NEBuilder HiFi DNA Assembly with a complementary oligonucleotide containing the random barcode sequence (Extended Data Table 4). The barcoded plasmid pool was transformed into MegaX DH10B T1R Electrocomp cells (Invitrogen, C6400-03), and colony-forming units were quantified to confirm >16,000-fold plasmid coverage. The sgRNA library was designed using CRISPick^34^. Oligonucleotide pools containing sgRNA sequences and compatible overhangs were cloned into the barcoded LentiCRISPRv2 backbone following BsmBI-v2 digestion and T4 DNA ligation. The final sgRNA-barcode library was transformed as above, achieving >10,000-fold sgRNA coverage. For validation experiments, individual sgRNAs were cloned into LentiCRISPRv2 (Addgene #52961) or TLCV2 (Addgene #87360).

### Lentivirus production

Lentiviral particles were produced in HEK293 cells using Lipofectamine 3000 (Thermo Fisher Scientific) in Opti-MEM medium (Gibco). Transfer plasmids were co-transfected with psPAX2 (Addgene #12260) and pCAG-VSV-G (Addgene #35616). Transfection medium was replaced after 24 h, and viral supernatant was collected 24 h later. Supernatants were filtered and concentrated by PEG precipitation according to the manufacturer’s protocol (Abcam, ab102538). Viral pellets were resuspended in mTeSR medium (STEMCELL Technologies), aliquoted, and stored at -80 °C until use.

### ORIGAMI CRISPR screen and barcode filtering

iPSCs were transduced with the ORIGAMI sgRNA-barcode library at a multiplicity of infection of 0.2, maintaining approximately 1,000 cells per sgRNA. Transduction was performed by centrifugation at 910 x g for 1 h at 31 °C in the presence of 8 μg/mL polybrene. Cells were washed three times with PBS after transduction. Forty-eight hours later, cells were selected with puromycin (0.3 μg/mL) for 4 days. For each biological replicate and time point, a minimum of 1.5 million cells was collected for genomic DNA extraction using the ISOLATE II Genomic DNA Kit (Bioline). sgRNA-barcode regions were amplified using primers listed in Extended Data Table 4, and PCR products were purified using the ISOLATE II PCR and Gel Kit (Bioline). Next-generation sequencing was performed by Novogene. Reads were processed in R using the ShortRead package^35^ to generate an sgRNA-barcode count matrix.

Barcode-level molecular identifier counts were examined independently for each sgRNA. Sequencing data from biological replicates were first combined, and the UMI counts associated with each unique barcode belonging to a given sgRNA were collected. For each sgRNA *g*, the mean barcode abundance (*μ*_g_) and standard deviation (*σ*_g_) were calculated across all associated barcodes:

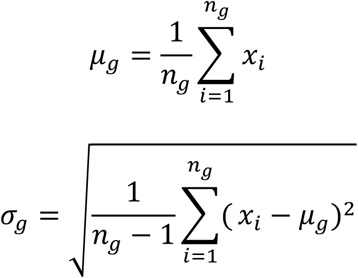

Where *x_i_* is the UMI count for barcode *i*, and *n*_g_ is the total number of barcodes associated with sgRNA *g*. Barcode-level outliers were defined as any barcode satisfying

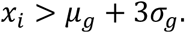

and were considered likely to represent founder-driven clonal expansion rather than reproducible sgRNA enrichment and thus excluded from downstream analysis. Following filtering barcode counts were summed to generate sgRNA-level counts, which were used as input for differential enrichment analysis using MAGeCK-MLE^36^. The effects of bar code filtering were assessed by visual inspection of barcode mean-variance relationships before and after filtering (Extended Data Figure 1C), confirming that removal of extreme barcode outliers reduced founder-driven variability while preserving the overall distribution of barcode abundances.

### Stem cell maintenance

All lines used in these studies were approved by the University of Sydney Human Ethics Committee (HREC 2020/417). Cell lines used in these studies included SCTi003-A(Stemcell Technologies, #200-0511) and ATCC-BXS0116 (ATCC ACS-1030). Induced pluripotent stem cells were maintained on Matrigel (Merck, CLS354277) coated plates and fed with MTSER or MTSER-plus (Stemcell Technologies, 85857 or 100-1130, respectively). Cells were routinely passaged using RelesR (Stemcell technologies 100-0483). Lines were karyotyped at the beginning of experiments and not used beyond 20 passages.

### Organoid generation

Cerebral organoids were generated using the STEMdiff Cerebral Organoid Kit (STEMCELL Technologies, #08570) following established protocols ^37^. Briefly, 50-70% confluent iPSCs were dissociated to single cells, and 9,000 cells were plated per well into ultra-low-attachment U-bottom plates in embryoid body formation medium supplemented with 10 μM ROCK inhibitor on DIV0. Medium was changed every other day until DIV5, when embryoid bodies were transferred to induction medium. On DIV7, organoids were embedded in Matrigel and cultured in expansion medium until DIV10, after which they were maintained in maturation medium for the remainder of differentiation. Organoids were transferred to low-attachment plates and cultured on an orbital shaker from DIV12 onward.

### Immunostaining

Organoids were fixed in 4% paraformaldehyde for 2 h at room temperature or overnight at 4 °C. Samples were washed in PBS and cryoprotected by sequential equilibration in 15% sucrose and 30% sucrose for 24 h each at 4 °C. Organoids were embedded in Tissue Freezing Medium (Leica), snap frozen, cryosectioned, and stored at -80 °C until staining. For immunostaining, slides were equilibrated to room temperature, washed in PBS, permeabilized in PBS containing 0.3% Triton X-100 for 15 min, and blocked for 1 h in PBS containing 5% normal goat serum, 1% bovine serum albumin, 0.3 M glycine, and 0.05% Triton X-100. Primary antibodies were diluted in blocking buffer and incubated overnight at 4 °C. Samples were washed three times in PBS containing 0.05% Triton X-100 and incubated with secondary antibodies diluted in blocking buffer at room temperature, followed by three additional washes. Antibodies are listed in Extended Data Table 5. The same staining protocol was used for immunocytochemistry of cells plated in 96-well format.

### Image acquisition and quantification

Brightfield images were acquired using an inverted Zeiss Axio microscope and Zeiss acquisition software (Zeiss, Zen V3.13.109). Organoid area was quantified in Fiji/ImageJ^38^. Fluorescence images were acquired using the Opera Phenix Plus High-Content Imaging System (Revvity), and image analysis was performed in Harmony software (Revvity). For nuclear marker quantification, nuclei were segmented using Hoechst/DAPI signal and marker-positive nuclei were identified using intensity thresholds applied consistently across control and experimental groups. For radial intensity analyses, line-scan or region-based measurements were performed from the ventricular edge toward the outer organoid layer over a distance of 200 μm. Area-under-the-curve values were calculated from intensity profiles and compared across genotypes.

### RT-qPCR

RNA was isolated using the ISOLATE II RNA Mini Kit (Bioline), and cDNA was synthesized using the iScript Select cDNA Synthesis Kit (Bio-Rad). RT-qPCR was performed using SYBR Select Master Mix (Thermo Fisher Scientific) on a QuantStudio 7 Real-Time PCR System (Thermo Fisher Scientific). Relative expression was calculated using the ΔΔCt method after normalization to a reference gene (*GAPDH*). Primer sequences are listed in Extended Data table 4.

### Molecular cloning

The coding sequence for human FBXW11 (Extended Data Table 4) was synthesized as a gBlock (Integrated DNA Technologies) and cloned into a lentiviral expression vector (Addgene #61425) using NEBuilder HiFi DNA Assembly. A matched empty-vector control was assembled in parallel. For proximity-labelling experiments, the FBXW11 lentiviral expression construct was modified by NEBuilder HiFi DNA Assembly to introduce miniTurboID at either the N terminus or C terminus of FBXW11. miniTurboID was amplified from Addgene #107172 after removal of the nuclear localization signal. Additional oligonucleotides and cloning details are provided in Extended Data Table 4.

### Single-Cell RNA-Seq preparation

Organoids were collected and diced using sterile scalpels in 15mm dishes (10 organoids were collected per genotype, per differentiation). These were dissociated to single cell suspensions using the Papain Dissociation System (Worthington Biochemical Corporation) according to standard protocols^39^. For cell hashing, single cell suspensions were incubated in Cell Staining Buffer (420201 BioLegend) and Fc Blocking reagent (FcX, BioLegend) for 10 mins at 4C. Cells were washed in staining buffer, and incubated in the same buffer using 0.5 μg of respective cell hashing antibodies (Extended Data Table 5) for 20 mins at 4C. Cells were washed twice with staining buffer, counted, and combined (4000 cells/hashtag, 16000 cells/reaction) before loading onto 10X Chromium Chip. Libraries were prepared according to the manufacturer’s protocol using the Chromium GEM-X Single Cell 3 Feature Barcode Kit.

### Quantifying Single-Cell Expression

Using Cell Ranger v8.8.0, each pooled transcriptomic library was aligned to reference GRCh38 (GENCODE v44) using “count” only quantifying exonic mapped reads. Pool A and Pool B were supplied with feature of HTO sequences.

### Single Cell Demultiplexing

Count matrices for gene expression (from Cell Ranger) and HTO and CITE-seq antibodies (from CITE-seq-Count) were imported into R using the Seurat v4. For each pool, HTO counts were normalised with centred log ratio (CLR) transformation, then assigned sample classification using Seurat’s HTODemux command (positive quantile=0.96, kfunc=”clara”, seed=11, and nsamples=2, Extended Data Fig. 3c). 2 hashtags corresponding to Control C1 and FBXW11 KO C1. Whilst for group 2: Control C2 and FBXW11 KO.

### Normalization, Filtering and Celltype Clustering

Only cells containing demultiplexed HTO and RNA-expressed were used for downstream analysis. Other filters included removing doublets HTO classification, cells with mitochondrial percentage above 15, cells with less than 400 and greater than 1000 genes expressed, and cells with less than 1000 and greater than 6500 UMIs quantified. Integration across Pool A and B was performed using Harmony. RNA count data was scaled and normalized using Seurat “LogNormalise” and function and the top 2000 variable features found using “vst” selection method were used to scale and centre features. Principal component analysis was run on the logged counts for 50 dimensions. An Elbow plot was used to select the number of dimensions (n dimensions = 30) for dimensionality reduction via Uniform Manifold Approximation and Projection (UMAP). To remove the effect of IgH genes on clustering and cell-type assignment IgH genes were removed. Clusters assigned to the Leiden algorithm were tested at a range of resolutions, visualized with clustree and a resolution of 0.4 chosen.

### TurboID proximity labelling and mass spectrometry

Proximity labelling was performed as described previously ^40^ with minor modifications. Cells stably expressing FBXW11-miniTurboID were supplemented with biotin for 10 min at 500 μM, washed five times in PBS, and lysed in RIPA buffer containing 1% Triton X-100, 0.1% SDS, 1% sodium deoxycholate, 150 mM NaCl, and 10 mM Tris-HCl pH 7.5, supplemented with cOmplete EDTA-free protease inhibitors (Roche, #11836170001). Biotinylated proteins were captured using streptavidin-conjugated magnetic beads (Thermo Fisher Scientific, #11205D) overnight at 4 °C. Beads were washed five times in 50 mM Tris-HCl and twice in 2 M urea prepared in 50 mM Tris-HCl pH 7.5. Bound proteins were digested on-bead with trypsin for 1 h at 25 °C with shaking, reduced with DTT to 4 mM for 30 min, and alkylated with iodoacetamide to 10 mM for 45 min in the dark. Proteins were further digested overnight with trypsin at 25 °C. Peptides were acidified with 1% trifluoroacetic acid, desalted using HLB columns (Waters, #186000383), dried by vacuum centrifugation, and resuspended in 3% acetonitrile/0.1% trifluoroacetic acid. Samples were sonicated for 5 min and 5 μL was analysed on a Q Exactive HF-X mass spectrometer (Thermo Fisher Scientific). Proteomic data were analysed using Spectronaut.

### Co-immunoprecipitation and immunoblotting

For co-immunoprecipitation, cells expressing HA-tagged FBXW11 were lysed under non-denaturing conditions using Pierce IP Lysis buffer (Thermo Fisher Scientific, #87787). Lysates were cleared by centrifugation and incubated with anti-HA antibody (abcam, ab9110) or an IgG control overnight at 4C in a rotator. After incubation, magnetic beads (Pierce Magnetic G beads) were added and incubated for 4 hours using a rotator at 4C. Immunoprecipitates were then washed once with TBS-tween (0.2 %) and eluted in Laemmli sample buffer. Input, unbound, and immunoprecipitated fractions were resolved by SDS-PAGE and transferred to membranes for immunoblotting. Membranes were probed with antibodies against FBXW11, HA, and CTNNB1/β-catenin as indicated in Extended Data Table 4. Signals were detected using LICOR.

### BrdU incorporation and cell-cycle analysis

Cells were incubated with 10 μM 5-bromo-2’-deoxyuridine (BrdU) in mTeSR Plus for 1 h. Cells were washed twice with PBS and dissociated to single cells using Accutase (STEMCELL Technologies, #07922). Cells were fixed in ice-cold ethanol for 30 min and washed twice in FACS buffer containing PBS, 2% bovine serum albumin, and 0.05% Tween-20. DNA was denatured with 2 M HCl for 1 h at room temperature. Cells were washed twice and incubated with anti-BrdU antibody (Supplementary Table 5) for 1 h at room temperature, followed by secondary antibody and Hoechst staining for 30 min. Cells were washed and analysed on a Cytek Aurora flow cytometer (Cytek Biosciences). Cell-cycle phase distribution was quantified based on BrdU incorporation and DNA content.

### FBXW11 Variant assembly and classification

*FBXW11* missense variants were assembled from two sources: ClinVar records for the gene and published de novo variants reported in patients with neurodevelopmental disorder with jaw, eye and digital anomalies. Variants were classified as pathogenic or de novo when reported as pathogenic or likely pathogenic in ClinVar or as de novo in a published clinical report, and as variants of uncertain significance when so classified in ClinVar. Because several substitutions are reported in more than one source and at more than one numbering convention, the pathogenic set was reduced to unique substitutions at the level of position plus substituted amino acid, giving 10 unique pathogenic substitutions at 9 residues.

Fourteen ClinVar records classified as pathogenic for FBXW11 are large chromosomal copy-number events (5q35 microduplication; Hunter-McAlpine craniosynostosis) rather than within-gene variants. These carry no protein position, span 11.95 to 31.56 Mb, contain 124 to 217 protein-coding genes each, and all four of the FBXW11-containing events also contain NSD1. They were excluded from all missense analyses. One missense variant of uncertain significance (MANE p.Pro63Leu) falls inside the MANE-specific insertion and has no UniProt equivalent, giving a comparator set of 83 rather than 84.

### Structural analysis

Distances were computed from the mmCIF deposition of PDB 6WNX, a 2.5 A structure of FBXW11 in complex with a doubly phosphorylated beta-catenin degron peptide (See Extended Data Table 6). The asymmetric unit contains three copies of the complex; protein chains A, D and G were paired with peptide chains C, F and I respectively.

Two features of the deposition are load-bearing and were handled explicitly. First, the degron phosphoserines are deposited as the non-standard component SEP, which many structure parsers discard by default; because the phosphate groups constitute the recognition chemistry of a phosphodegron, all SEP atoms were retained. Second, distances were minimised across all three crystallographic copies rather than computed from a single copy.

For each residue, the reported distance is the minimum heavy-atom to heavy-atom distance between that residue and any residue of the bound peptide, taken as the minimum over the three protein-peptide pairs. Hydrogens were not considered.

Deposited numbering in 6WNX corresponds to UniProt isoform Q9UKB1-2 (508 residues) and maps to the canonical 542-residue sequence by an offset of +113. This offset was verified by amino-acid identity at all 410 modelled positions of chain A. Across the three copies the structure resolves canonical residues 114 to 524 (chain A 114-523, chain D 115-522, chain G 114-524); because distances are minimised over all three protein-peptide pairs, the union is the relevant coverage. Residues outside that span have no distance and were excluded rather than imputed. Contact residues were defined as those within 4.5 A of the bound peptide, giving 24 residues; a stricter 4.0 A threshold gives a nested subset of 22.

### TOPFlash Luciferase assay

Stem cells were transfected with the β-catenin-responsive TOPFlash luciferase reporter (Addgene #12456 ^19^) using Lipofectamine 3000 (Thermo Fisher Scientific). Medium was replaced the following day, and cells were treated with CHIR99021 at the indicated concentrations or DMSO vehicle control for 24 h. Cells were counted for normalization and lysed using the Steady-Glo Luciferase Assay System (Promega). Luciferase activity was normalized to cell number and expressed relative to the corresponding control condition.

## Data availability

The single-cell 10x 3’ RNA-seq raw sequencing data the associated gene x count matrix generated for this study has been deposited at the European Nucleotide Archive (ENA) under accession number E-MTAB-17543.

## Use of generative AI and AI-assisted technologies in the manuscript preparation process

During the preparation of this manuscript, the authors used ChatGPT (OpenAI) and Claude (Anthropic) to assist with language editing, improving clarity and readability, and refining the presentation of the manuscript. These tools were not used to generate primary data or to independently perform or interpret the scientific analyses. All AI-assisted content was reviewed and edited by the authors, who take full responsibility for the final content of the manuscript.

## Acknowledgments

GGN was supported by the NHMRC (GNT2020532, GNT1185002, GNT1107514, GNT1158164, GNT1158165, GNT1046090, GNT1111940) a kind donation from Dr. John and Anne Chong. CLM was supported by a kind donation from Dr. John and Anne Chong. The authors would like to acknowledge the Sydney Cytometry Facility and the Sydney Microscopy Facility for technical assistance.

## Author contributions

CLM, GGN designed the study. CLM, ST, and GGN developed ORIGAMI. CLM, HK, and RW helped with single cell data analysis and interpretation. CLM, ST, MT, DB and TYD helped with organoid generation and immunohistochemistry. CLM, TM, and GGN helped with *Drosophila* studies and interpretation. CLM and GGN drafted the main manuscript, and all other authors helped edit.

## Supplementary Materials

Supplementary Table 1. Complete sgRNA Library.

Supplementary Table 2. Differentially expressed genes in FBXW11 deficient iPSCs.

Supplementary Table 3. Number of Cells per cell type and condition (FBXW11 KO and Control).

Supplementary Table 4. Oligonucleotides for cloning, library preparation, and RT-qPCR.

Supplementary Table 5. Antibodies.

Supplementary Table 6. FBXW11 clinical variants distances.

## Extended Data Figures

**Extended Data Fig. 1.**
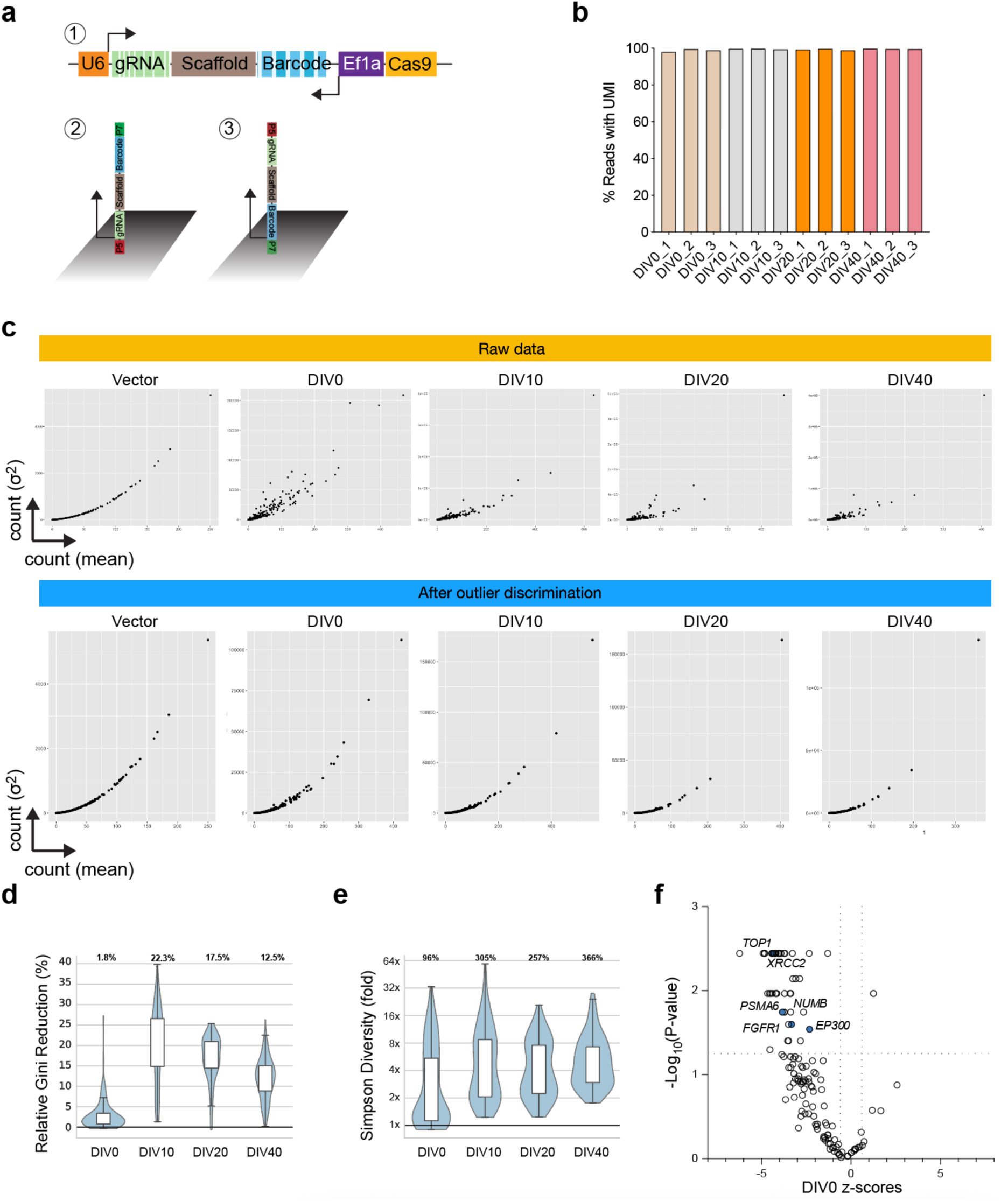
Additional data on ORIGAMI CRISPR screen. **a,** Schematic showing CRISPR targeting design. (1) Simplified vector design with random barcode after sgRNA scaffold. Constant regions preceding the gRNA sequence and Ef1alpha promoter are used in library generation for next generation sequencing. Using paired end readings (2) the P5 adapter is used to capture the sgRNA sequence and the (3) P7 adapter is used to read the barcode. **b,** Percent of sgRNA reads with detected barcode across the screen. **c,** Mean sgRNA count and variance before barcode correction shows expected parabolic relationship, which deteriorates as organoids differentiate. After outlier discrimination, extreme barcode-driven variance is reduced while the expected mean–variance relationship is preserved. **d,** Relative reduction in Gini inequality after barcode-level outlier discrimination across DIV0, DIV10, DIV20, and DIV40. **e,** Relative improvement in Simpson diversity after barcode-level outlier discrimination across DIV0, DIV10, DIV20, and DIV40. Values indicate the median fold increase at each time point. Increased diversity after filtering indicates reduced dominance of individual barcode outliers. **f,** Baseline screen behaviour of positive-control and cell-fitness genes at DIV0. Several positive control fitness-associated genes were depleted at the stem cell stage including: *FGFR1*, *EP300*, *PSMA6*, *XRCC2*, *TOP1*, and *NUMB*.

**Extended Data Fig. 2.**
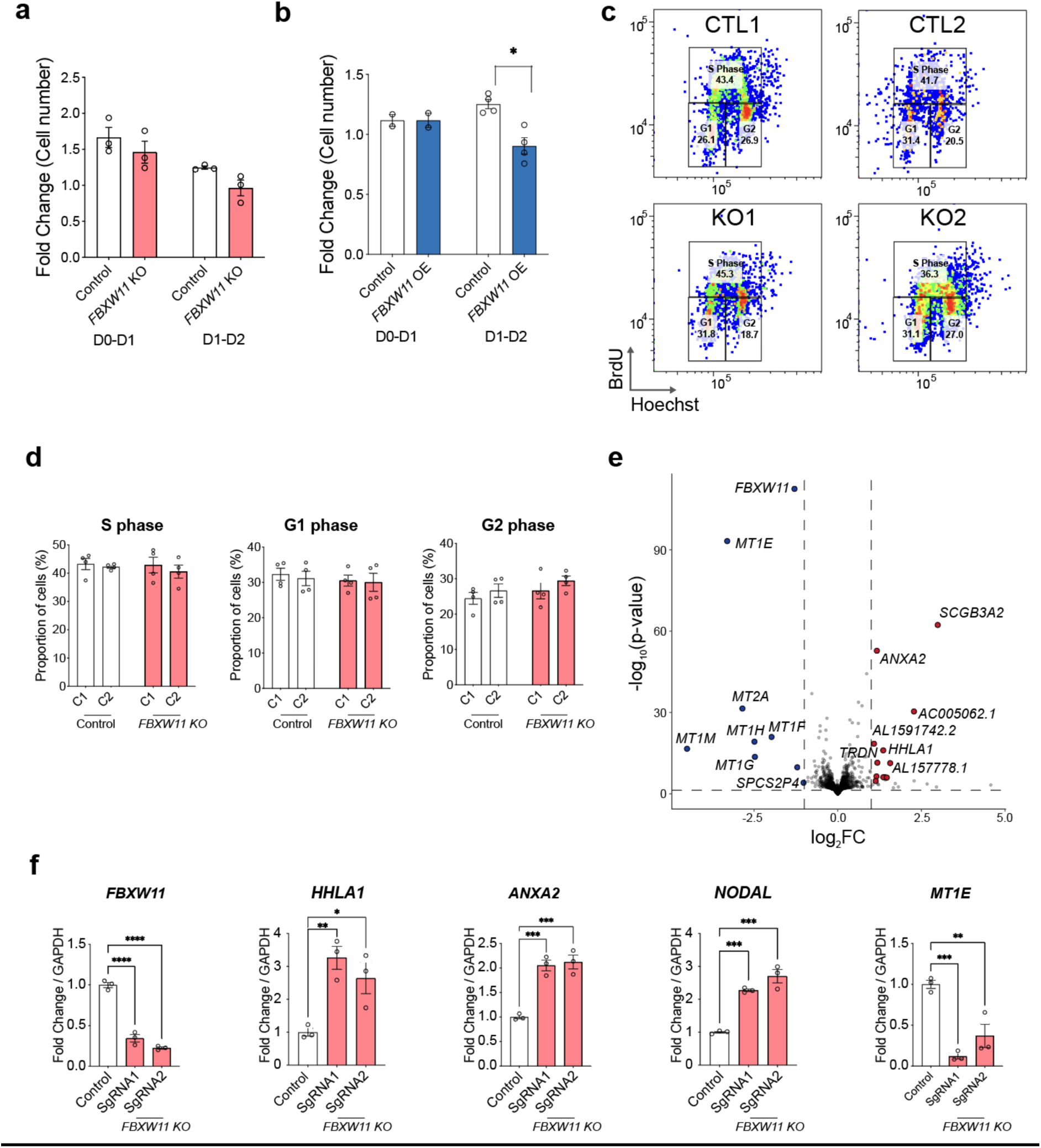
Additional phenotypes in response to *FBXW11* expression. **a,** Fold change differences across days in culture for *FBXW11* KOs and controls. N=3 experimental repeats. **b,** Fold change differences across days in culture for *FBXW11* overexpressing cells and controls. 2-way ANOVA. *, P<0.05. N=2-4 experimental repeats. **c,** Representative flow cytometry plots after BrdU pulse to discriminate cell cycle in stem cells with *FBXW11* KO or control. **d,** Quantification of (C), n = 4 experimental replicates. **e,** Volcano plot showing differentially expressed genes as determined by RNA-sequencing in *FBXW11* KO cells vs. control. n = 3 experimental replicates. **f,** Individual RT-qPCR validation of top differentially expressed genes identified by RNA-Seq. n = 3 experimental replicates.

**Extended Data Fig. 3.**
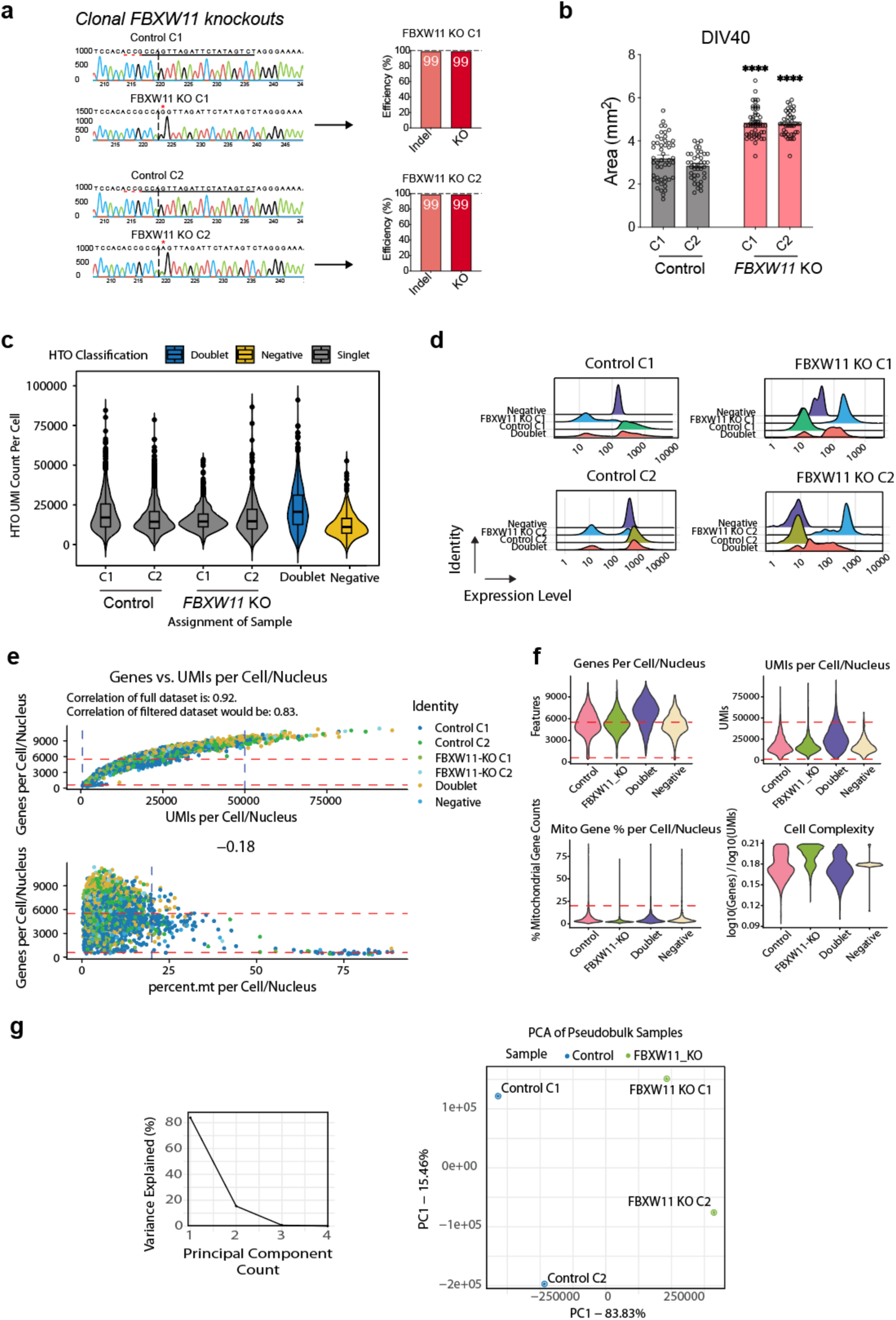
Single RNA-Seq Supporting Data. **a,** Validation of clonal *FBXW11* knockout iPSC lines used for downstream single-cell RNA-seq. Representative Sanger sequencing traces are shown for two independent control clones and two independent FBXW11 KO clones. ICE analysis estimated indel and knockout efficiencies. **b,** Quantification of DIV40 cerebral organoid area in control and FBXW11 KO lines. Each dot represents an individual organoid across n =3 experimental replicates. Bars indicate mean ± SEM. Statistical significance was determined relative to controls; ****, P < 0.0001. **c,** Hashtag oligonucleotide (HTO) UMI counts per cell/nucleus following sample multiplexing. Cells/nuclei were classified as singlets, doublets, or negative based on HTO enrichment, enabling demultiplexing of control and *FBXW11* KO samples prior to downstream analysis. **d,** Representative HTO expression distributions used for sample assignment. Ridge plots show HTO signal across each sample identity, doublet, and negative classification, supporting separation of control and *FBXW11* KO libraries after multiplexing. **e,** Single-cell/nucleus RNA-seq quality-control metrics. Scatter plots show the relationship between detected genes and UMI counts, and between detected genes and mitochondrial read percentage. Red dashed lines indicate filtering thresholds applied to remove low-quality cells/nuclei and high-mitochondrial-content events. Correlation values before and after filtering are shown. **f,** Violin plots summarising quality-control metrics across retained sample classes, including detected genes per cell/nucleus, UMIs per cell/nucleus, mitochondrial gene percentage, and cell complexity. These metrics were used to assess dataset quality prior to integration and clustering. **g,** Dimensionality reduction quality control for pseudobulk samples. The elbow plot shows variance explained across principal components, and PCA of pseudobulked samples separates control and *FBXW11* KO samples along the major principal components while retaining clone-level structure.

**Extended Data Fig. 4.**
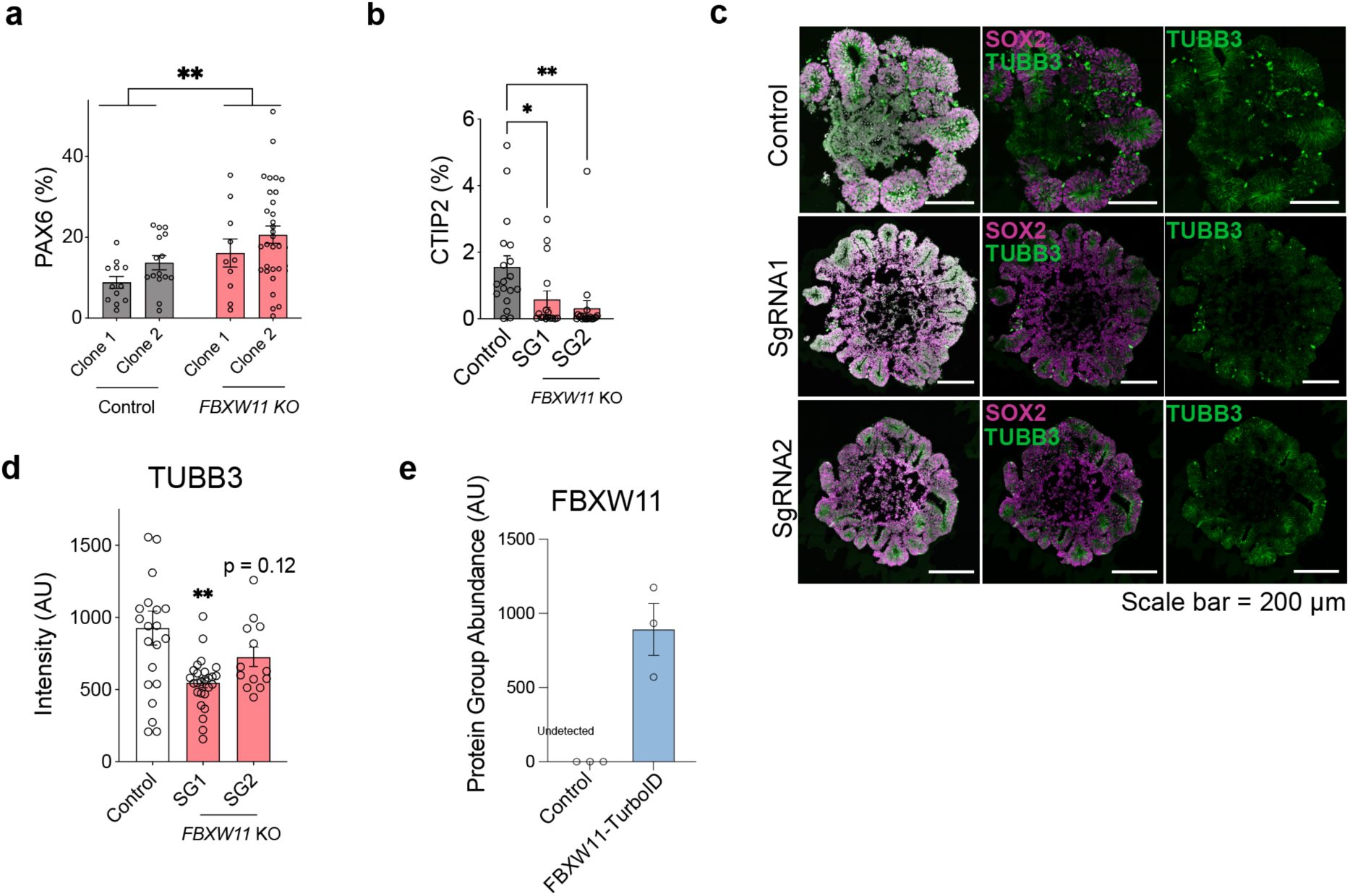
Additional characterisations. **a,** Quantification of PAX6-positive nuclei across Control and *FBXW11* KO organoids DIV40. Each dot represents an individual organoid across n = 3 independent differentiations. Statistical significance was determined by two-way ANOVA; **P < 0.001. **b,** Quantification of CTIP2-positive nuclei across Control and *FBXW11* KO DIV40 organoids. Each dot represents an individual organoid across n = 3 independent differentiations. Statistical significance was determined by one-way ANOVA Dunnett post hoc test; *P<0.05; **P < 0.001. **c,** Representative immunofluorescence images of DIV10 cerebral organoids stained for SOX2, and TUBB3. **d,** Quantification of TUBB3-positive regions across Control and *FBXW11* KO DIV40 organoids. Each dot represents an individual organoid across n = 3 independent differentiations. Statistical significance was determined by one-way ANOVA Dunnett post hoc test; **P < 0.001. **e,** Protein group abundance detected using mass spectrometry in FBXW11-TurboID expressing stem cells vs. control (empty vector).

